# Dynamic protein phosphorylation shapes mitochondrial catalysis, import, and architecture

**DOI:** 10.64898/2026.09.28.755151

**Authors:** Andrew J. Smith, Patrick Forny, Sean W. Rogers, Merima Forny, David J. Pagliarini

## Abstract

Phosphorylation is a pervasive mitochondrial protein post-translational modification, yet the extent of its influence over mitochondrial functions remains unclear. To identify dynamic mitochondrial phosphoisoforms, we profiled the phosphoproteome of AML12 cells following individual genetic perturbations of 10 resident mitochondrial phosphatases. Guided by these results, we investigated the functional implications of phosphorylation on mitochondrial processes related to catalysis, import, and morphology. We find that phosphorylation of adenylate kinase 2 suppresses its nucleotide binding and catalytic activity. Elevated phosphorylation proximal to the mitochondrial targeting sequence of branched chain ketoacid dehydrogenase kinase disrupts its import and processing, resulting in enhanced catabolism of branched chain amino acids. Most notably, we discover extensive phosphorylation on proteins involved in mitochondrial cristae architecture, including multiple members of the MICOS complex and ATP synthase, upon silencing the phosphatase PGAM5. Collectively, our work connects reversible phosphorylation and resident phosphatases to the regulation of diverse mitochondrial processes.

**HIGHLIGHTS:**

- Perturbation of 10 mitochondrial resident phosphatases maps dynamic phosphosites.
- AK2 S151 phosphorylation tunes catalysis by augmenting adenine nucleotide binding.
- MTS-proximal BCKDK phosphorylation impairs protein import and gates BCAA catabolism.
- MIC19 and ATP5I phosphorylation remodels cristae and modulates bioenergetics.

## INTRODUCTION

Reversible protein phosphorylation is among the most pervasive and versatile mechanisms of cellular regulation and is controlled reciprocally by kinases and phosphatases. Phosphorylation acts as a molecular switch that reshapes protein conformation, stability, localization, and molecular interactions, tuning processes that range from signal transduction to metabolism. The many mechanisms through which a single modified residue can act, and the aberrations that follow when this control fails, underscore both the physiological importance of phosphorylation and the difficulty of inferring its function from occurrence alone.

Nearly all cells rely on mitochondria for their core metabolic needs and must respond to acute shifts in nutrient availability and energy demands. As such, it is essential that cells can appropriately maintain the content, function, and activity of these organelles. As a rapid and reversible post-translational modification (PTM), phosphorylation is well-suited to help calibrate these demands. The discovery of pyruvate dehydrogenase regulation by phosphorylation almost 60 years ago cemented the importance of this PTM in mitochondrial biology^1^. Continued technological advances in mass spectrometry have since cataloged thousands of mitochondrial phosphosites that are reproducibly dynamic across physiological states and disease, a subset of which have defined functional consequences^2–4^. The great majority of these sites occur at low stoichiometry and have no ascribed function, leaving open whether they are regulatory or largely incidental^5,6^. Distinguishing consequential phosphorylation from the bystander majority is therefore a central obstacle to understanding this modification in mitochondria.

Despite this uncertainty, the presence of multiple resident phosphatases suggests that protein dephosphorylation may be broadly important for calibrating mitochondrial activities. These phosphatases, many of which are conserved across eukaryotic species, are poorly characterized and span distinct catalytic domains and mitochondrial compartments^7^. Their disruption can produce specific and often severe phenotypes; most strikingly, loss of the matrix/OMM phosphatase PPTC7 causes profound metabolic dysfunction and perinatal lethality in mice through dysregulated mitochondrial content^8,9^. That cells devote conserved machinery to removing these modifications suggests that mitochondrial phosphorylation may be broadly regulatory — or at least consequential — even where individual sites occur at low stoichiometry.

To define which mitochondrial proteins carry functionally consequential phosphorylation, we individually depleted ten resident mitochondrial phosphatases and quantified resulting changes across the mitochondrial phosphoproteome. This revealed a large set of dynamically regulated phosphosites, preferentially located within intrinsically disordered regions. To nominate phosphosites likely to be functional, we applied three complementary analytical frames, each prioritizing candidates by a distinct principle: structural proximity, which flags sites near catalytic or ligand-binding residues as candidate direct effectors of enzyme activity; positional enrichment, which flags phosphorylation clustered at defined sequence landmarks such as targeting and processing signals; and phosphorylation density, which flags individual proteins and multiprotein complexes carrying a heavy phosphorylation load. Pursuing one or more candidates from each frame, we uncovered regulatory roles spanning enzyme catalysis, protein import and maturation, and the assembly of the complexes that shape cristae, while leaving many further candidates for future study. Collectively, these findings establish mitochondrial phosphorylation as a widespread, dynamic, and functionally consequential modifier of core mitochondrial processes.

## RESULTS

### The mitochondrial phosphoproteome is dynamic

Although nearly 90% of the mitochondrial proteome carries at least one reported phosphoisoform, the overwhelming majority of these sites lack any published function (Fig. S1A). We reasoned that bona fide regulatory phosphorylation sites would be subject to modification by resident mitochondrial phosphatases (Fig. 1A). We therefore used CRISPRi to silence the expression of 10 such phosphatases (Fig. 1B) in AML12 mouse hepatocytes expressing dCas9-KRAB-MeCP2. Medium- and Heavy-SILAC cells received control or phosphatase-targeting sgRNAs, respectively, and were selected, expanded, and combined 1:1 prior to mitochondrial isolation and proteomic/phosphoproteomic profiling. Phosphopeptides exhibited both shared and uniquely regulated clusters (Fig. 1C), underscoring the breadth and dynamism of mitochondrial phosphorylation.

**Figure 1.**
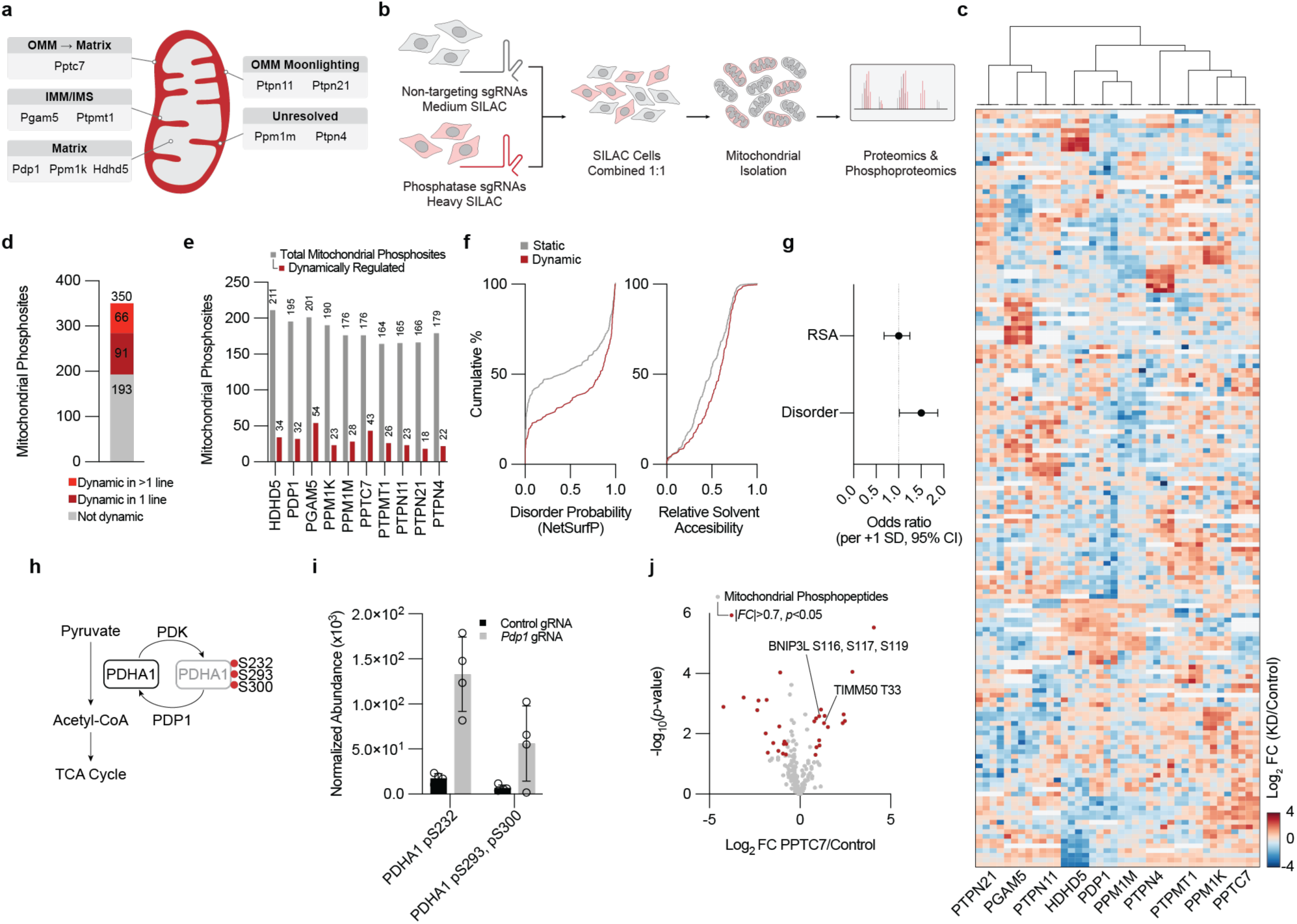
CRISPRi perturbation of resident mitochondrial phosphatases maps the dynamic mitochondrial phosphoproteome. (A) Schematic of the ten resident mitochondrial phosphatases targeted for knockdown, grouped by sub-mitochondrial localization. (B) Experimental design: AML12 cells expressing dCas9-KRAB-MeCP2 were transduced with non-targeting control (Medium-SILAC) or phosphatase-targeting (Heavy-SILAC) sgRNAs, selected, expanded, combined 1:1, and profiled by proteomics and phosphoproteomics after mitochondrial isolation. (C) Hierarchical clustering of dynamically regulated phosphosites across the ten knockdown lines. |Log2 FC| ≥ 1 and Benjamini–Hochberg (BH)-adjusted p < 0.05 (Welch t-test on protein-normalized phosphopeptide values; n = 4 SILAC replicates per line). (D) Number of identified mitochondrial phosphosites (350 total): not dynamic (193), dynamic in one line (91), and dynamic in more than one line (66). (E) Identified (grey) and dynamically regulated (red) mitochondrial phosphosites per knockdown line. (F) Predicted disorder probability and relative solvent accessibility (RSA) for dynamically regulated (n = 157) and static (n = 192) mitochondrial phosphosites (one of the 350 identified sites lacked a structural model and was excluded from structural analyses). Disorder 0.83 vs. 0.43, p = 2×10⁻⁴, RSA 0.59 vs. 0.49, p = 0.008; two-sided Mann–Whitney U. Relative solvent accessibility from Shrake–Rupley SASA on AlphaFold models and disorder probability from NetSurfP predictions over the MitoCarta3.0 proteome. (G) Odds ratios (points) with 95% confidence intervals (bars) for association with dynamic status, from multivariable logistic regression with continuous predictors standardized to unit standard deviation (OR per +1 SD); dashed line indicates OR = 1 (no association). (H) Diagram of pyruvate entry into the TCA cycle: PDK-mediated phosphorylation of PDHA1 (S232, S293, S300) inhibits the complex, and PDP1 dephosphorylates and reactivates it. (I) Normalized abundance of PDHA1 pS232 and pS293/pS300 in control versus Pdp1-knockdown cells. (J) Volcano plot of mitochondrial phosphopeptides in PPTC7-knockdown versus control; known PPTC7 substrates BNIP3L (S116, S117, S119) and TIMM50 (T33) are elevated (|log2FC| > 0.7, p < 0.05, Welch t-test on non-normalized phosphopeptide values; n = 4 SILAC replicates per line).

Across the dataset we identified 350 mitochondrial phosphosites, 157 of which (∼45%) were dynamic — 91 in a single line and 66 in multiple — capturing both uniquely and commonly regulated sites (Fig. 1D). Although each knockdown identified a similar number of sites overall (164–211), only a small fraction changed in any given line (Fig. 1E), meaning the full dynamic repertoire emerged only across the panel. Examining the occurrence of phosphorylation across the mitochondrial proteome, we found it to be unevenly distributed across proteins, with a median of 1 phosphoform per protein (Fig. S1B). Overall, we found that phosphorylation occurred predominantly in loop regions (Fig. S1C) but that dynamic sites were much more frequently associated with disordered regions than were static sites (median predicted disorder 0.83 vs. 0.43; Fig. 1F). They were also modestly more solvent-exposed (median relative solvent accessibility (RSA) 0.59 vs. 0.49), but a joint logistic-regression model attributed this entirely to disorder: disorder alone predicted dynamic status (odds ratio (OR) 1.55 per standard deviation (SD)), whereas accessibility, after adjustment, did not (Fig. 1G). Dynamic mitochondrial phosphoregulation is therefore linked to intrinsic disorder rather than solvent accessibility per se.

To gauge the validity of our data, we next asked whether it recovered phosphosites with well-established phosphatase regulation. Entry of pyruvate into the TCA cycle is governed by reversible phosphorylation of pyruvate dehydrogenase (PDHA1) at S232, S293, and S300. PDK-mediated phosphorylation inhibits the complex and blocks pyruvate oxidation, while PDP1 removes these modifications to restore activity (Fig. 1H). Consistent with the loss of its cognate phosphatase, all three PDHA1 sites were elevated in PDP1-knockdown cells relative to controls (Fig. 1I). The same sentinel behavior held for PPTC7, whose knockdown increased phosphorylation of its known targets BNIP3L/NIX (S116, S117, S119) and TIMM50 (T33) (Fig. 1J). Thus, our approach faithfully recaptured established regulatory phosphatase–substrate relationships, and inspires confidence that our resulting dataset could reveal additional, as-yet-uncharacterized phosphoregulatory dynamics.

### Phosphorylation is an inhibitory modifier of Adenylate Kinase 2

A phosphosite lying within an enzyme’s active site can alter catalysis directly, by reshaping the geometry or electrostatics of substrate binding^10,11^, whereas a regulatory site acting from a distance implies allosteric control. To nominate phosphosites of these classes, we ranked dynamically regulated enzyme phosphosites by their proximity to core catalytic residues. Of the 350 identified mitochondrial phosphosites, 113 lay on enzymes (72 distinct proteins), and 53 of these were dynamically regulated (40 proteins). Of these 53 sites, 49 belonged to enzymes with well-annotated active site residues, thus allowing us to measure the three-dimensional distance between the phospho-acceptor hydroxyl to the nearest atom of an annotated catalytic or ligand-binding residue (Fig. 2A). Because a phosphate extends ∼4 Å beyond the acceptor oxygen, we classified the six sites within 8 Å (phosphate reach and van der Waals contact) as direct-effect candidates positioned to contact the active site, and the remaining 43 as candidate regulators acting at a distance.

**Figure 2.**
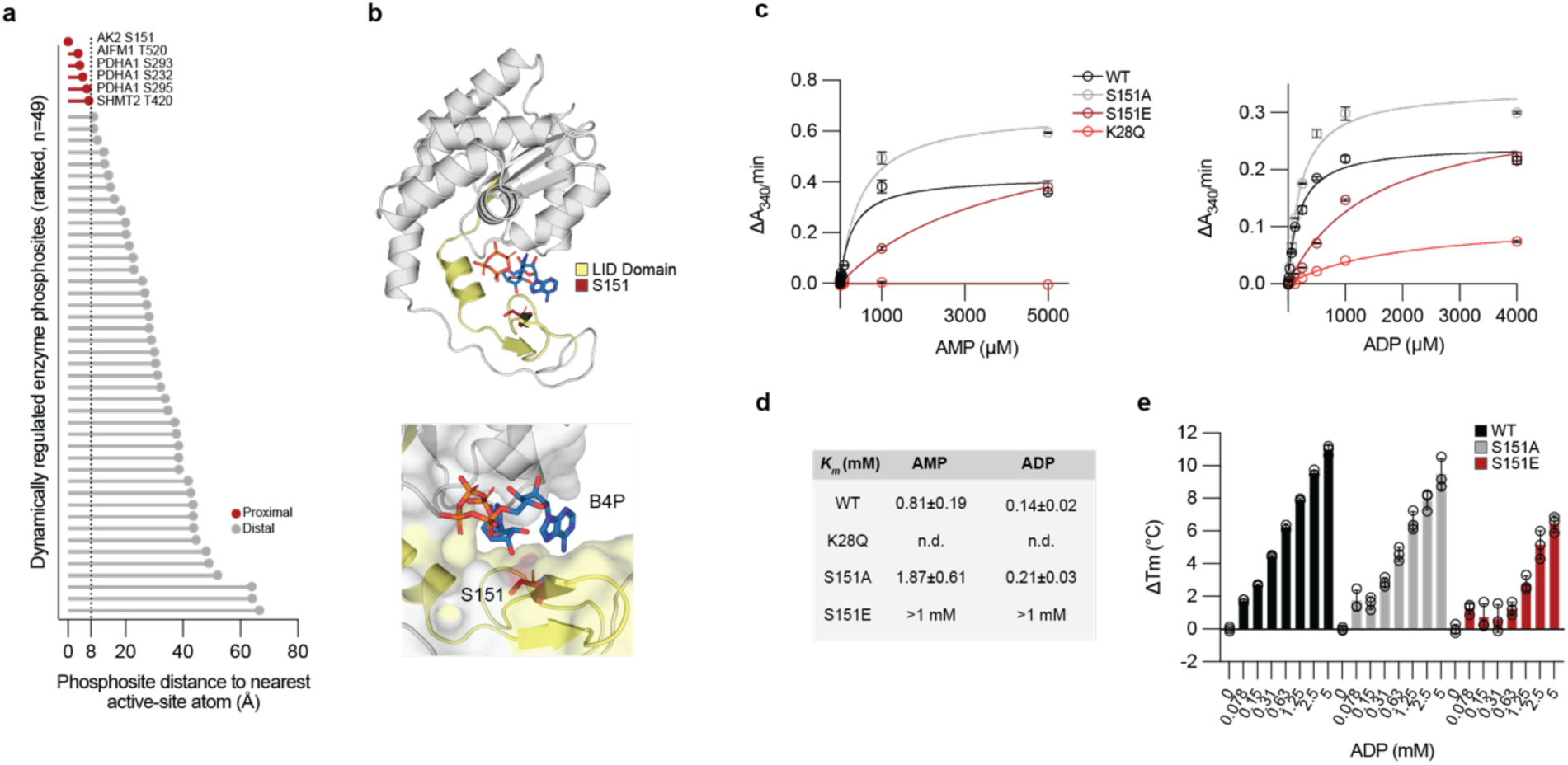
Phosphorylation of a LID-domain site directly inhibits adenylate kinase 2 (AK2). (A) Regulated phosphosites ranked by proximity to a catalytic active site; AK2 S151 ranked highest. Sites ranked by proximity to catalytic active sites using structural coordinates and active-site annotations (AlphaFold/AlphaFill models, and UniProt active/binding-site annotations). (B) Structure of AK2 bound to P1,P4-di(adenosine-5′)-tetraphosphate (B4P), S151 in the LID domain (PDB: 2C9Y). (C) Steady-state kinetics (forward, AMP varied; reverse, ADP varied) for WT, K28Q, S151A, S151E. Michaelis–Menten nonlinear regression; errors = fit SE. n = 2 preparations. (D) Km table for steady-state kinetics. (E) Differential scanning fluorimetry (ΔTm) across ADP concentrations for WT, S151A, S151E.

This partition recovered established phosphoregulatory sites in both classes, supporting its validity. Among the 43 distal sites was S79 of acetyl-CoA carboxylase (ACACA), a canonical allosteric regulatory site, alongside uncharacterized candidates including MTHFD2 S149 — down-regulated across three knockdown lines (PDP1, PGAM5, PTPN21) — and ACADL S193, specifically elevated upon HDHD5 depletion (Fig. S2C, D). The six direct-effect candidates were AK2 S151 (0 Å, ATP), AIFM1 T520 (3.5 Å, NAD⁺), the PDHA1 sites S232/S293/S295 (4–6.6 Å, thiamine diphosphate/Mg²⁺), and SHMT2 T420 (7.2 Å, folate). Reassuringly, this set re-identified the known regulatory sites of PDHA1 and surfaced a novel proximal site, S295, elevated upon PTPN11 loss (Fig. S2B), as well as AIFM1 T520, positioned 3.5 Å from the NAD⁺-coordinating residue W195 (Fig. S2A). We selected the highest-ranked candidate — AK2 S151, on the intermembrane-space adenylate kinase AK2 — for mechanistic dissection.

To understand how S151 phosphorylation might affect AK2, we examined its position in the structure of AK2 bound to the bisubstrate analog P1,P4-di(adenosine-5′)-tetraphosphate (B4P) (Fig. 2B). S151 lies in the LID domain — the mobile subdomain that closes over the active site during catalysis and supplies positively charged residues that engage substrate and stabilize the transition state^12–14^. In the closed, substrate-bound conformation S151 sits within the nucleotide-binding cleft, adjacent to the bound substrate, such that a phosphomonoester at this position would introduce a negative charge into the pocket and electrostatically disfavor nucleotide binding. To test this, we purified recombinant wild-type (WT), phosphodead (S151A), phosphomimetic (S151E), and P-loop catalytic-dead (K28Q) AK2 (Fig. S2E) and measured steady-state kinetics of the forward (AMP + ATP → 2 ADP) and reverse (2 ADP → AMP + ATP) reactions (Fig. 2C). K28Q was inactive in both directions, confirming that the signal reflects genuine AK2 chemistry. S151A modestly increased the Michaelis constant (∼2-fold; AMP 1.9 mM vs. 0.805 mM, ADP 0.21 mM vs. 0.14 mM for WT), whereas S151E apparently weakened substrate affinity so severely that its Km could not be reliably determined within the assayed range in either direction (Km > 1 mM for AMP and ADP) (Fig. 2D). That both reaction directions were affected identifies residue 151 as a determinant of nucleotide affinity. We corroborated this by differential scanning fluorimetry, measuring ligand-induced thermal stabilization across ADP concentrations (Fig. 2E). WT and S151A were progressively stabilized by ADP (ΔTm ∼11 and ∼9 °C), whereas S151E showed markedly reduced stabilization (ΔTm ≤ ∼6 °C) that emerged only at high ADP, mirroring its elevated Km. Together, these results establish S151 as a phosphorylation-sensitive determinant of AK2 nucleotide binding, and show how a single modification positioned within an active site can directly tune a mitochondrial enzyme’s activity.

### MTS-proximal phosphorylation influences protein processing and metabolic outputs

Mitochondrial protein import and maturation are increasingly recognized as points of regulatory control, and phosphorylation of precursor proteins within the targeting presequence or the adjacent segment processed by MPP and MIPEP/XPNPEP3 peptidases offers one plausible mechanism^15–17^. Whether such phosphorylation is positionally concentrated near the targeting sequence, and whether these sites are dynamically regulated, remains undefined at proteome scale. We reasoned that a regulatory role for phosphorylation at the mitochondrial N-terminus would manifest as a positional bias in phosphosite occurrence, with dynamic sites concentrating there.

To test this while controlling for the short, compositionally skewed presequences, we mapped each of the 350 identified phosphosites in precursor coordinates relative to the MitoFates-predicted MPP cleavage site and normalized occurrence to local phosphoacceptor content (phosphosites per 100 S/T/Y; Fig. 3A). Of the 350 phosphosites, 142 (59 dynamic) mapped to 85 MTS-containing proteins. A continuous positional profile (11-residue sliding window, ±5) showed phosphorylation depleting within the MTS and rising sharply immediately after the cleavage site (pooled mean 0.43 per 100 S/T/Y for all sites, 0.18 for dynamic sites; Fig. 3A). We defined the MTS-proximal region as the first 30 mature residues — the window of peak dynamic phosphosite density and the segment engaged by the import and processing machinery. Dynamic phosphosites were 4.0-fold enriched in this proximal zone relative to the distal remainder (>30; p = 9.8 × 10⁻⁶), whereas the MTS itself was phospho-depleted (Fig. 3B).

**Figure 3.**
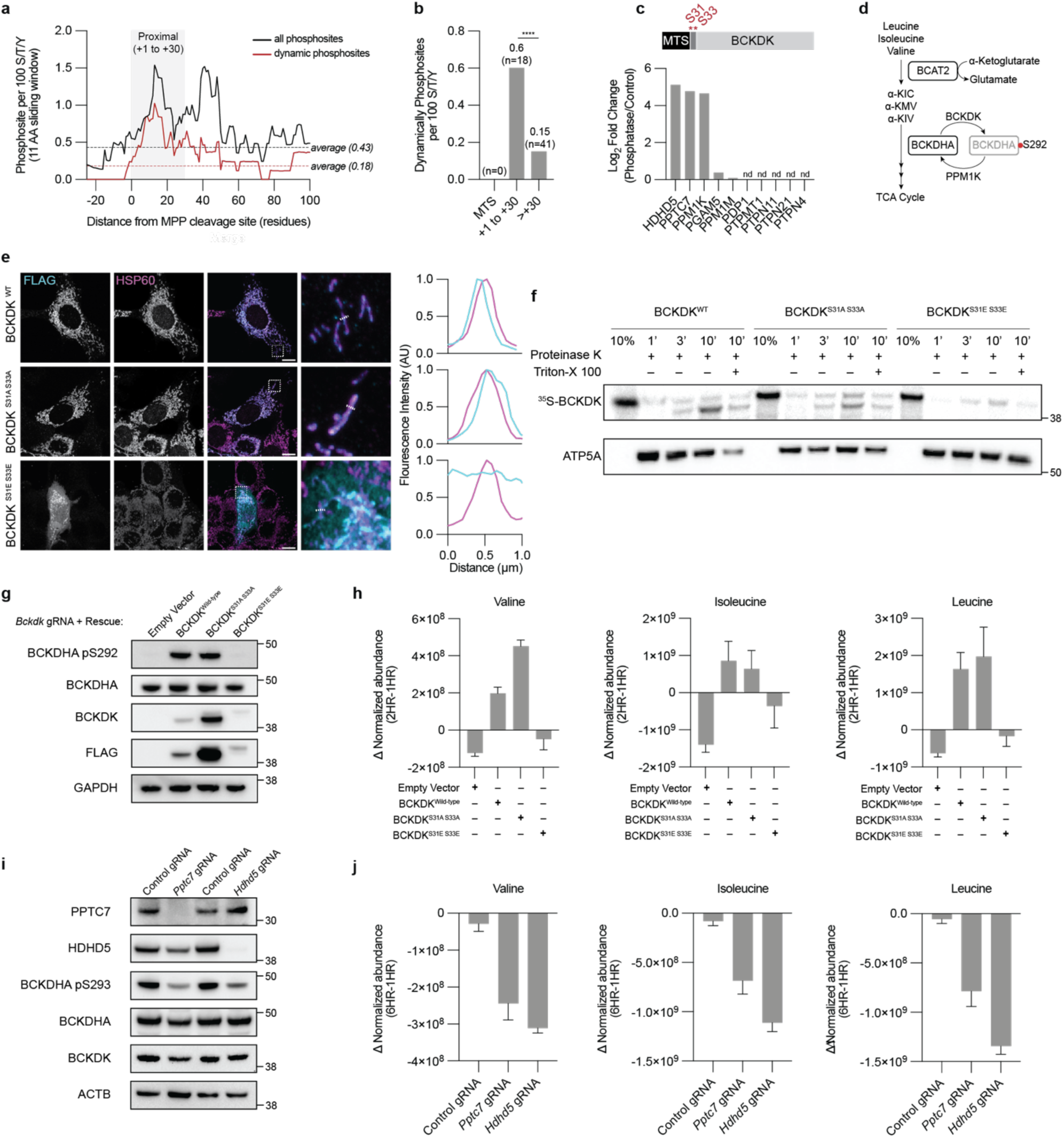
Positional phosphorylation adjacent to targeting sequences couples mitochondrial import and maturation to metabolic output. (A) Total and dynamic phosphosite occurrence normalized to S/T/Y content relative to MPP cleavage site. (B) Dynamic phosphosites per 100 S/T/Y in the MTS, proximal (+1 to +30), and distal (>30) amino acids from MPP cleavage site. Dynamic phosphorylation was 4.0-fold enriched in the proximal zone relative to the distal remainder (P = 9.8 × 10⁻⁶, two-sided Fisher’s exact). (C) Log2 fold change (knockdown/control) of BCKDK S31/S33 across 10 lines; induction in HDHD5 (≈5.1), PPTC7 (≈4.8), PPM1K (≈4.6), nd = not detected. Adjusted p (S31/S33): PPTC7 6.9 × 10⁻⁵, PPM1K 1.4 × 10⁻⁴, HDHD5 5.3 × 10⁻⁴; n = 4 per channel. (D) Schematic of BCAA catabolism. BCKDK phosphorylates/inhibits BCKDHA (S293/S303), PPM1K reactivates it; red circles = BCKDHA phosphorylation. (E) Immunofluorescence of FLAG-tagged (cyan) WT, S31A/S33A, and S31E/S33E BCKDK in AML12 cells. Mitochondria are stained with HSP60 (magenta). Representative line-profiles for both channels are measured across regions indicated in the inset. Scale bar = 10 µm. (F) Mitochondrial import and processing assay of [^35^S]-labelled BCKDK proteins in HEK-293T cells. ATP5A is used as a loading control. Representative of N = 3 experiments. (G) Immunoblots of BCKDK-KO cells reconstituted with EV, WT, S31A/S33A, or S31E/S33E. (H) Pulse-chase experiment measuring [^13^C]-labelled Valine, Isoleucine, and Leucine in BCKDK rescue cell lines. Data is differential normalized abundance (Area/Protein/Arginine×1^12^) between 2 and 1 hours (Data is ± SEM, N=3 replicates per group). (I) Immunoblots of PPTC7-KO and HDHD5-KO cells with control gRNA transduced cells. (J) Pulse-chase experiment measuring [^13^C]-labelled Valine, Isoleucine, and Leucine in control, PPTC7-KO and HDHD5-KO cells. Data is differential normalized abundance (Area/Protein/Arginine×1^12^) between 6 and 1 hours (Data is ± SEM, N=3 replicates per group).

BCKDK S31/S33 was the strongest proximal event and was elevated across multiple lines (HDHD5, PPTC7, PPM1K; Fig. 3C), suggesting that its regulation may be responsive to more general metabolic perturbations. In contrast, other proximal sites were line-specific. For example, an N-terminal ETFA phosphopeptide (S21/T22/S32/T37) was enriched only upon PTPN4 knockdown (Fig. S3A), and both ISCU S15/S30 (Fig. S3B) and NAXE S43 (Fig. S3C) were only elevated upon PPTC7 knockdown, consistent with our prior reports^9^. Given its relevance across multiple conditions, we focused on BCKDK S31/S33 for functional dissection.

BCKDK is the principal negative regulator of BCAA catabolism, phosphorylating and inactivating the BCKDH complex to gate entry of leucine, isoleucine, and valine into oxidative metabolism (Fig. 3D). To begin to understand the consequence of BCKDK S31/S33 phosphorylation, we first tested for an effect on localization. We expressed WT, non-phosphorylatable (S31A/S33A), and phosphomimetic (S31E/S33E) BCKDK and found that S31E/S33E was sufficient to impair full mitochondrial localization (Fig. 3E). We next measured in vitro mitochondrial import of recombinant [^35^S]-labeled BCKDK. All constructs were imported, as shown by time-dependent accumulation of proteinase K-protected species (Fig. 3F); however, S31E/S33E was imported more slowly than WT and S31A/S33A, and its processing was disrupted, with only full-length (i.e., uncleaved) S31E/S33E accumulating in mitochondria. Because MTS removal by MPP is required for mature-protein stability, these data indicate that phosphorylation proximal to the BCKDK targeting sequence dampens its import, and that dephosphorylation after translocation is required for processing, a paradigm consistent with our previous work^9^.

We reasoned that if S31/S33 phosphorylation blocks productive import, the phosphomimetic mutant should phenocopy BCKDK loss. In BCKDK-knockout cells reconstituted with empty vector, WT, S31A/S33A, or S31E/S33E, only WT and S31A/S33A drove BCKDHA S293 phosphorylation whereas S31E/S33E and empty vector did not, with BCKDHA abundance unchanged (Fig. 3G). Accordingly, in a pulse-chase experiment, leucine, isoleucine, and valine accumulated in WT and S31A/S33A cells but were consumed in empty-vector and S31E/S33E cells (Fig. 3H). To ask whether this regulation operates endogenously, we examined knockouts of PPTC7 and HDHD5, two phosphatases that oppose BCKDK S31/S33 phosphorylation (Fig. 3C). Both reduced BCKDHA S293 phosphorylation relative to control despite unchanged BCKDHA abundance (Fig. 3I) and both depleted branched-chain amino acids more rapidly in a pulse-chase (Fig. 3J), phenocopying the phosphomimetic mutant. Together, these data reveal that disruptive phosphorylation immediately downstream of the BCKDK targeting sequence impairs its import and processing, thereby relieving BCKDH inhibition and accelerating BCAA catabolism. Collectively, these data demonstrate the feasibility of a regulatory paradigm that couples the efficiency of a protein’s import and maturation to a defined metabolic output.

### Densely phosphorylated proteins alter cristae architecture

Phosphorylation can also act cumulatively, where modifications distributed across the many subunits of a shared assembly constitutes a coordinate regulatory input^18^. To assess how phosphorylation is distributed across mitochondrial proteins, we ranked all 190 phosphorylated mitochondrial proteins (350 identified phosphosites; 106 of these proteins carrying 157 dynamically regulated sites) by phosphosite count normalized to protein length. This ranking revealed a strongly right-skewed distribution (Fig. 4A, B): a small number of proteins were densely phosphorylated while the majority carried sparse modification (median 0.0045 phosphosites per residue), with density falling steeply with rank (Fig. 4A). The most densely phosphorylated proteins were not the large, abundant metabolic enzymes but a set of comparatively small regulators of mitochondrial dynamics, apoptosis, and cristae architecture — including BNIP3L, TIMM8A1, ATP5I, BCL2L13, CYB5B, MTFR1L, MFF, MIC27, and MIC19. The same trend held among dynamically regulated sites, again topped by regulators of mitochondrial dynamics and structure (MTFR1L, TIMM8A1, MIC27, BNIP3L, BAD) rather than by bulk metabolic machinery (Fig. 4B).

**Figure 4.**
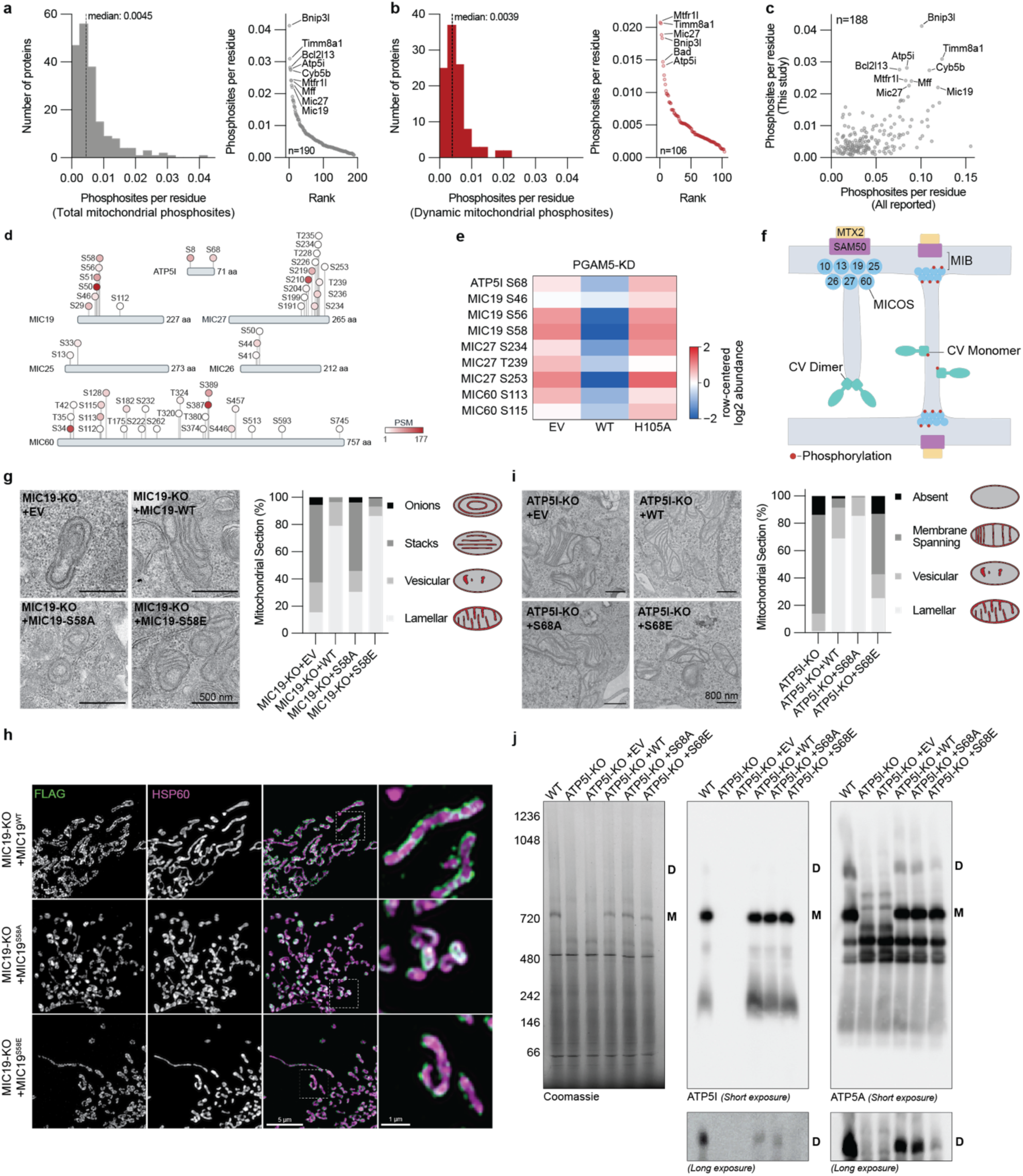
Dense phosphorylation of cristae regulators converges on MIC19 and ATP5I to control cristae architecture. (A) Length-normalized phosphosite density (left), and ranked (right) for all identified mitochondrial phosphosites (350 sites, 190 proteins). (B) Length-normalized phosphosite density (left), and ranked (right) the dynamically regulated subset (157 sites, 106 proteins; dynamic = protein-normalized |log2FC| ≥ 1 and BH-adjusted p < 0.05, Welch t-test). Selected top-ranked proteins labeled. (C) Length-normalized phosphosite density in this study versus the union of all reported mitochondrial phosphosites (dbPTM + EPSD), for the 188 proteins detected in both (Spearman ρ = 0.45, p = 9.4 × 10⁻¹¹); shared top-ranked proteins labeled. (D) Diagram of all identified phosphosites on proteins involved in cristae formation processes. Scale bar = number of peptide spectral matches (PSM) per phosphopeptide. (E) Row-normalized log2 abundance heatmap of cristae-formation phosphosites in PGAM5-KD rescue cells (EV, WT, H105A). (F) Schematic: MICOS-and outer membrane proteins SAM50 and MTX2 form the mitochondrial intermembrane space bridging complex (MIB) at crista junctions. ATP synthase dimers promote cristae tip formation, whereas monomers do not. Phosphorylation of both complexes influences cristae formation. (G) TEM of MIC19-KO rescues classified lamellar/vesicular/stacks/onions, with proportions; S58A fails to rescue, WT and S58E restore normal cristae. n = 142 (KO+EV), 138 (KO+WT), 131 (KO+S58A), and 130 (KO+S58E) mitochondrial sections scored across 2 independent experiments. (H) High-resolution confocal (FLAG, green; HSP60, magenta) of MIC19-KO+WT/S58A/S58E; WT and S58E form discrete puncta and rescue morphology, S58A is diffuse. (I) TEM of ATP5I-KO rescues classified lamellar/vesicular/membrane-spanning/absent, with proportions; WT and S68A rescue, S68E does not fully. n = 94 (KO), 106 (KO+WT), 90 (KO+S68A), and 91 (KO+S68E) mitochondrial sections scored across 2 independent experiments. (J) Blue-native PAGE Coomassie (loading control) and immunoblots for ATP5I and ATP5A (short/long exposure) showing monomers (M) and dimers (D); dimers are markedly reduced with S68E.

To test whether this distribution is a feature of our dataset specifically or of the mitochondrial phosphoproteome generally, we compared our per-protein density with that computed identically from the union of all previously reported mitochondrial phosphosites (Fig. 4C), and found this trend holds across all previously reported datasets. Dense phosphorylation of mitochondrial proteins is therefore a reproducible feature of the phosphoproteome rather than an artifact of a single study. To ask if this dense phosphorylation is directed toward specific mitochondrial functions, we weighted each of the 828 detected mitochondrial proteins by its number of dynamically regulated phosphosites and tested MitoCarta3.0-defined pathways for enrichment of this weighted signal against the rest of the proteome. This identified multiple regulatory programs of which the MICOS complex was the most densely phosphorylated (Fig. S4A).

Within the MICOS/cristae-formation program, phosphopeptide isoform analysis identified 50 phosphosites across the entire dataset (Fig. 4D). Restricting this set to phosphosites with confident site localization and reproducible quantification (detected in at least three of four replicates per line), 24 phosphosites were quantified with high confidence across the five detected MICOS subunits (MIC60, MIC19, MIC25, MIC27, MIC26) and ATP5I. PGAM5 knockdown accounted for the majority of the dynamic events detected (Fig. S4B). Re-expression of WT PGAM5, but not the catalytically dead H105A mutant, reversed the elevation of 12 highlighted sites (Fig. 4E, S4C), confirming their dependence on PGAM5 catalytic activity.

Among these sites, ATP5I S68 and MIC19 S58 were the most strongly PGAM5-dependent, and we selected them for mechanistic follow-up. To interpret these two sites structurally, we considered how their host complexes sculpt the inner membrane (Fig. 4F). The MICOS complex assembles at crista junctions — the narrow necks connecting cristae to the inner boundary membrane — generating negative membrane curvature, whereas rows of F_1_F_O_-ATP synthase dimers impose the strong positive curvature that forms cristae tips. The opposing activities of these two complexes are a fundamental determinant of cristae shape^19–22^. Because MIC19 is a core MICOS subunit and ATP5I an ATP synthase subunit, we asked whether phosphorylation of each site affects assembly of its respective complex.

To test this for MIC19, we reconstituted MIC19-knockout cells with WT, phosphodead (S58A), or phosphomimetic (S58E) MIC19. Immunoblotting confirmed loss of endogenous MIC19, comparable re-expression of each construct, and restoration of MIC60, whose levels track MICOS complex stability (Fig. S4D). Quantitative proteomic profiling of S58A- and S58E-reconstituted cells gave closely comparable protein-expression profiles (Fig. S4E), localizing any phenotypic difference to protein assembly rather than expression. By transmission electron microscopy, WT and S58E MIC19 restored normal lamellar cristae, whereas phospho-null S58A failed to rescue the knockout phenotype (Fig. 4G). High-resolution confocal microscopy reinforced this distinction: WT- and S58E-reconstituted MIC19 organized into discrete puncta and rescued the knockout’s gross morphological defects, whereas S58A remained diffuse and non-punctate (Fig. 4H).

We next applied the same design to ATP5I. ATP5I-knockout cells reconstituted with WT, S68A, or S68E ATP5I confirmed loss and re-expression by immunoblotting (Fig. S4F), and proteomic profiling showed that knockout-associated complex abundance defects were restored comparably by both phosphodead and phosphomimetic constructs (Fig. S4G), again localizing any phenotypic difference downstream of complex-member expression. By TEM, ATP5I-knockout mitochondria displayed a distinctive “membrane-spanning” cristae phenotype in which cristae fail to form discrete tips; WT and S68A ATP5I restored normal cristae, whereas phosphomimetic S68E did not (Fig. 4I). Blue-native PAGE localized this to complex assembly: monomeric and dimeric ATP synthase, absent in knockout and empty-vector cells, were restored by WT and S68A, while S68E supported markedly fewer dimers relative to monomers (Fig. 4J). All ATP5I constructs restored oxygen consumption in glucose (Fig. S4H), indicating that S68 phosphorylation acts through ATP synthase dimerization rather than through complex V catalytic activity.

Together, these results show that dynamic phosphorylation of the mitochondrial proteome is concentrated on a small set of structurally consequential proteins, and that two of its most heavily regulated members — MIC19 and ATP5I — each translate that phosphorylation into discrete effects on cristae architecture. Importantly, these data demonstrate that phosphorylation can positively or negatively regulate complex assembly, and suggests that these opposing effects may cooperatively promote cristae formation.

### Phosphorylation drives cristae remodeling and a bioenergetic growth advantage

To assess whether coordinated phosphorylation of MICOS/ATP synthase subunits is sufficient to remodel cristae architecture and cell physiology, we returned to PGAM5-knockdown mitochondria. We first assessed the submitochondrial topology of PGAM5 by proteinase K protection of isolated mitochondria (Fig. 5A) confirming an inner-membrane localization with the catalytic domain facing the IMS — positioning PGAM5 to act directly on the IMS-facing MICOS and ATP synthase subunits identified above. By TEM, PGAM5-knockdown mitochondria showed increased mitochondrial area and cristae density relative to control (Fig. 5B), consistent with enhanced MICOS/ATP-synthase-driven membrane remodeling.

**Figure 5.**
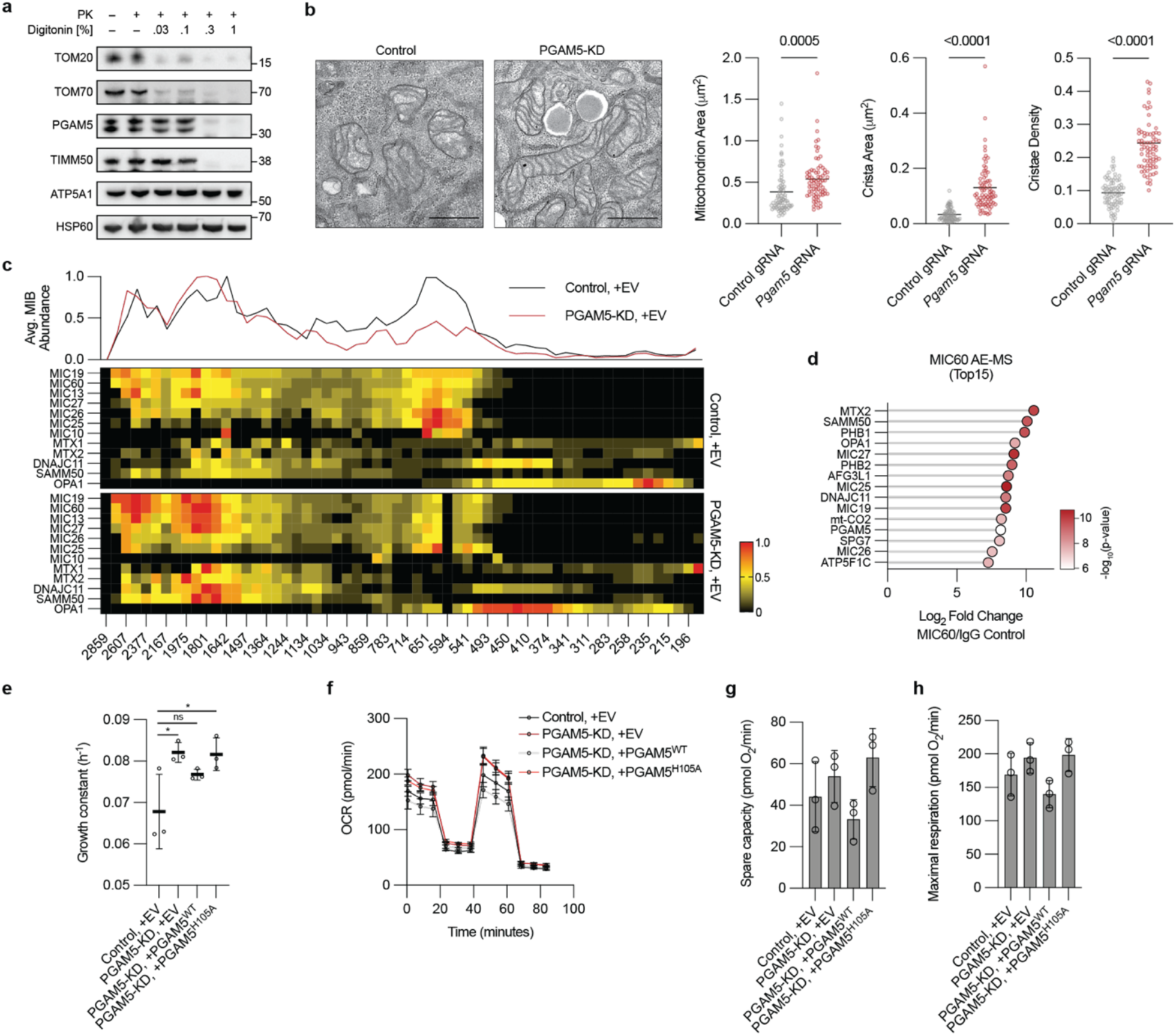
Cumulative phosphorylation remodels cristae architecture and cellular bioenergetics. (A) Proteinase K protection assay of isolated mitochondria immunoblotted for PGAM5, TIMM50 (IMM/IMS), TOM20/TOM70 (OMM), and ATP5A1/HSP60 (matrix); PGAM5’s profile aligns with TIMM50. Representative of N = 2 experiments. (B) TEM of control and PGAM5-KD mitochondria: representative micrographs and violin plots of mitochondrion area (Welch’s, P < 0.0005), cristae area (Welch’s, P < 0.0001), and cristae density (Welch’s, P < 0.0001). N = 74 mitochondria per condition. (C) Complexome profiling of MIB members and OPA1 (control vs PGAM5-KD), showing a shift toward the higher-MW MIB complex. (D) Log2 FC (MIC60/IgG), ranked top 15 significant MIC60-interacting proteins. Welch t-test, BH FDR, P < 0.05, n = 2. (E) Specific growth constant µ (h⁻¹) for control+EV, PGAM5-KD+EV, PGAM5-KD+WT, PGAM5-KD+H105A, over a 3–15 h exponential window. One-way ANOVA with Tukey’s post-hoc, *p<0.05, ns, not significant (n = 3 wells per condition, error bars are SEM). (F–H) OCR (pmol/min) in galactose for control+EV, PGAM5-KD+EV, PGAM5-KD+WT, PGAM5-KD+H105A, with spare capacity (G, maximal − basal respiration) and maximal respiration (H, max after FCCP − non-mitochondrial).

To first ask whether these enhanced cristae phenotypes simply reflect altered abundance of the affected proteins, we immunoblotted purified mitochondrial and cytosolic fractions from control and PGAM5-KD cells. PGAM5 depletion was confirmed in knockdown cells, but MIC19, MIC27, MIC60, ATP5I, and OPA1 levels were unchanged (Fig. S5A). Further, mitochondrial recruitment of DRP1 — a cytosolic GTPase previously reported to be activated by PGAM5-mediated dephosphorylation at the outer membrane — was likewise unaltered (Fig. S5A), consistent with PGAM5’s IMS-facing topology (Fig. 5A), together excluding altered protein stability, turnover, or DRP1-dependent fission as the basis for the phenotypes.

To examine MIB assembly directly, we performed complexome profiling of control and PGAM5-KD mitochondria. MICOS and additional MIB components (SAMM50, MTX1, MTX2, DNAJC11) redistributed toward the higher-molecular-weight MIB complex in PGAM5-KD (Fig. 5C), while the ratio of dimeric/oligomeric to monomeric complex V decreased (Fig. S5B) — consistent, respectively, with promotion of MICOS subunits into the mature MIB complex and with ATP5I phosphorylation restraining ATP synthase dimerization.

To determine whether PGAM5 physically engages this machinery, we performed crosslinking affinity-enrichment mass spectrometry of MIC60, the structural core of MICOS. PGAM5 was strongly and specifically enriched over IgG, alongside established MICOS-MIB interactors (MIC19, SAMM50, MTX2, OPA1, DNAJC11) (Fig. 5D), establishing PGAM5 as a bona fide component of the MIC60 interactome. Finally, to test the functional consequences of this axis, we assayed growth and respiration. In cell-growth assays, PGAM5-KD and H105A-rescued cells grew faster than control and WT-rescued cells in both glucose and galactose (Fig. 5E). In galactose, this was mirrored by a selective increase in spare respiratory capacity for the KD and H105A lines, whereas WT-rescued cells resembled control (Fig. 5F–H).

Together, coordinated phosphorylation of cristae-shaping complexes in PGAM5-knockdown cells tracks with increased MIB assembly, elevated cristae density, and a growth and respiratory advantage — establishing that loss of a single phosphatase is sufficient to convert site-level phosphorylation events into a whole-organelle phenotype.

## DISCUSSION

Mitochondrial protein phosphorylation has been recognized as physiologically consequential since the discovery of phosphorylation-dependent control of pyruvate dehydrogenase nearly 60 years ago^1^. Mass spectrometry-based approaches have since cataloged thousands of mitochondrial phosphosites^2–4^, the great majority without ascribed function^5,6^. Reasoning that phosphatase-targeted sites would likely hold functional value, we depleted ten resident mitochondrial phosphatases and monitored the resulting dynamic phosphoproteomic data to nominate candidates by three criteria: proximity to catalytic residues, positional enrichment near the MTS, and phosphorylation density across a shared complex. We found that each nominated a distinct, functionally validated mode of control: direct inhibition of AK2 catalysis, MTS-proximal gating of BCKDK import, and phospho-dependent assembly of MICOS and ATP synthase into mature cristae.

Constructing functional mitochondria already requires coordinating transcription, translation, import, and complex assembly. Our results suggest phosphorylation may add a further, likely faster-acting layer on top of this scaffold. In principle, because it requires no new protein synthesis, it can adjust composition (e.g., via import), catalytic output, and structural morphology on a timescale of seconds rather than the hours needed to remodel protein abundance. This may be particularly suited to cristae architecture, which requires cooperative assembly of many subunits rather than mere adjustment to individual subunit abundance^23^ — a property that would make it cumbersome to tune except by fast, reversible means, paralleling recent evidence that cardiolipin redistribution alone can rapidly reshape OPA1-dependent membrane curvature^24^.

For this to prove an accurate framework, multiple questions still need to be answered. First, are the sites we and others describe adventitious byproducts of a phosphorylation-permissive environment, or does a logic govern which sites recur? Second, and related, where does this phosphorylation originate — from resident kinases with broader substrate scope than currently appreciated^25,26^ (the PDKs, BCKDK, etc.,), from cytosolic kinases like PKA that have been reported to localize to mitochondria^27,28^ or to act on precursors before import (as our BCKDK data suggest), or non-enzymatically, as occurs in bacteria via reactive phosphodonors such as acetyl-phosphate^29^? Third, if these events are genuinely regulatory, do they operate through canonical, high-stoichiometry signal-transduction logic, or through a simpler mode potentially more consistent with mitochondria’s bacterial ancestry — perhaps acting through the same intrinsic disorder that was the strongest structural correlate of dynamic phosphorylation in our dataset, and that can drive switch-like conformational transitions in other disordered proteins^30^? Resolving these will require moving beyond steady-state snapshots to measure phosphorylation dynamics directly in response to meaningful biological perturbations.

Beyond the processes examined here, mitochondrial phosphorylation is increasingly implicated in disease: PINK1-mediated phosphorylation of ubiquitin and Parkin triggers mitophagy, and its loss causes Parkinson’s disease^31^; tau phosphorylation is linked to mitochondrial dysfunction in Alzheimer’s disease^32^; and cannabinoid signaling is reported to modulate memory via PKA-dependent phosphorylation of the complex I subunit NDUFS2^33^. Together with our data, these examples raise the possibility that altered mitochondrial phosphorylation may emerge as an underappreciated hallmark of mitochondrial dysfunction or as a means of organellar specialization^34–36^. If so, the phosphatases and kinases that set these marks could become attractive drug targets: we have previously shown that a small-molecule kinase inhibitor can be selectively directed to mitochondria to inhibit COQ8A^37^, and a fuller map of mitochondrial phospho-regulatory pathways could similarly inform mitochondria-targeted therapeutics.

More broadly, our data argue that reversible phosphorylation is a widespread and functionally impactful layer of mitochondrial regulation, acting through distinct mechanisms on catalysis, import, and architecture. Beyond the sites we pursued to mechanistic resolution, our dataset nominates many further candidates across all three prioritization frames that will serve as a rich foundation for further exploration.

## LIMITATIONS OF THE STUDY

Our screen is built around steady-state phosphorylation measured after an extended period of phosphatase depletion, and this design carries two related limitations. First, it captures the endpoint of chronic phosphatase loss rather than the real-time dynamics of phosphorylation. Second, because we perturbed the phosphatase side of the reaction, our approach is inherently biased toward sites under active phosphatase control; sites that turn over too quickly to accumulate under chronic depletion, would not be captured by this design. Similarly, our approach does not resolve phosphorylation stoichiometry at most sites in the dataset. Although fold-change magnitude suggests occupancy is high for a subset of sites, we cannot exclude that many of the phosphosites we describe as dynamic occur at overall low stoichiometry within the total cellular pool. This issue is compounded by emerging evidence that mitochondria within a single cell can form metabolically and functionally distinct subpopulations; if particular phosphorylation events are concentrated within one such subpopulation, the bulk biochemical measurements used here would systematically underestimate their true local abundance and functional significance. Finally, our functional experiments were performed in a single immortalized mouse hepatocyte line (AML12). While this system provided the throughput required for our needs, whether any individual regulatory relationship we describe generalizes to other tissues, cell types, or in vivo contexts remains to be established.

## MATERIALS AND METHODS

### KEY RESOURCES TABLE

#### Antibodies

| REAGENT or RESOURCE | SOURCE | IDENTIFIER |
| --- | --- | --- |
| Anti-ACTB | Abcam | ab8224 |
| Anti-MIC60/IMMT | Proteintech | 10179-1-AP |
| Anti-TOM20 | Cell Signaling Technology | 42406 |
| Anti-TIMM50 | Proteintech | 22229-1-AP |
| Anti-HSP60 | EnCor Biotech | CPCA-HSP60 |
| Anti-ATP5A | Proteintech | 14676-1-AP |
| Anti-ATP5I | Atlas Antibody | HPA035010 |
| Anti-FLAG | Cell Signaling Technology | 14793; D6W5B |
| Anti-BCKDK | Santa Cruz | sc-374424 |
| Anti-BCKDHA | Bethyl | A303790AT |
| Anti-BCKDHA pS293 | Bethyl | A304672AT |
| Anti-GAPDH | Cell Signaling Technology | 2118 |
| Anti-PPTC7 | Abcam | ab122659 |
| Anti-HDHD5 (CECR5) | Abcam | ab184185 |
| Anti-TOM70 | Proteintech | 14528-1-AP |
| Anti-PGAM5 | Thermo | PA5-57894 |
| Anti-MIC19 | Sigma | HPA042935 |
| Anti-MIC27 | Thermo | PA5-51427 |
| Anti-DRP1 | Cell Signaling Technology | 8570 |
| Anti-OPA1 | Cell Signaling Technology | 80471 |
| Anti-RFP | Proteintech | 6G6 |
| Rabbit IgG control | Thermo | 02-6102 |
| HRP-conjugated anti-rabbit IgG | Cell Signaling Technology | 7074 |
| HRP-conjugated anti-mouse IgG | Cell Signaling Technology | 7076 |
| Goat anti-Rabbit IgG, Alexa Fluor 488 | Invitrogen | A11008 |
| Goat anti-Chicken IgY, Alexa Fluor 647 | Invitrogen | A32933 |
| Protein A Dynabeads | Thermo | 10001D |

#### Bacterial and virus strains

| REAGENT or RESOURCE | SOURCE | IDENTIFIER |
| --- | --- | --- |
| E. coli Rosetta(DE3)pLysS | Sigma | 70956-3 |
| E. coli 5-alpha competent | New England Biolabs | C2987 |

#### Chemicals, peptides, and recombinant proteins

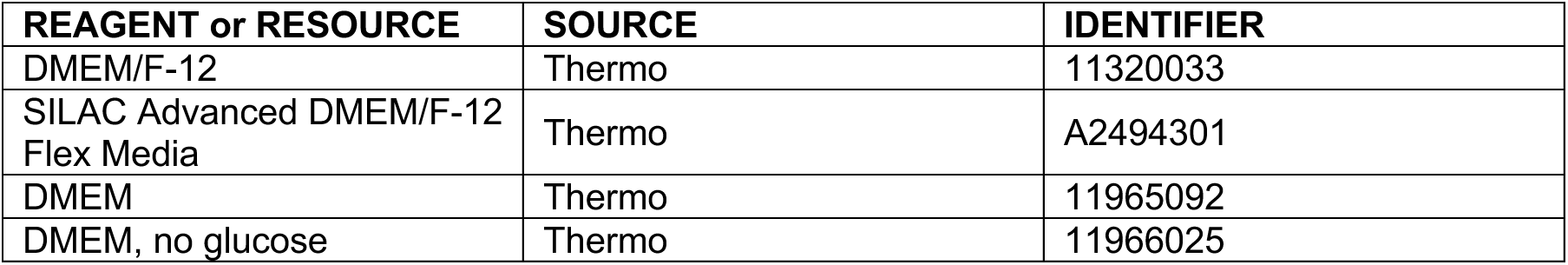
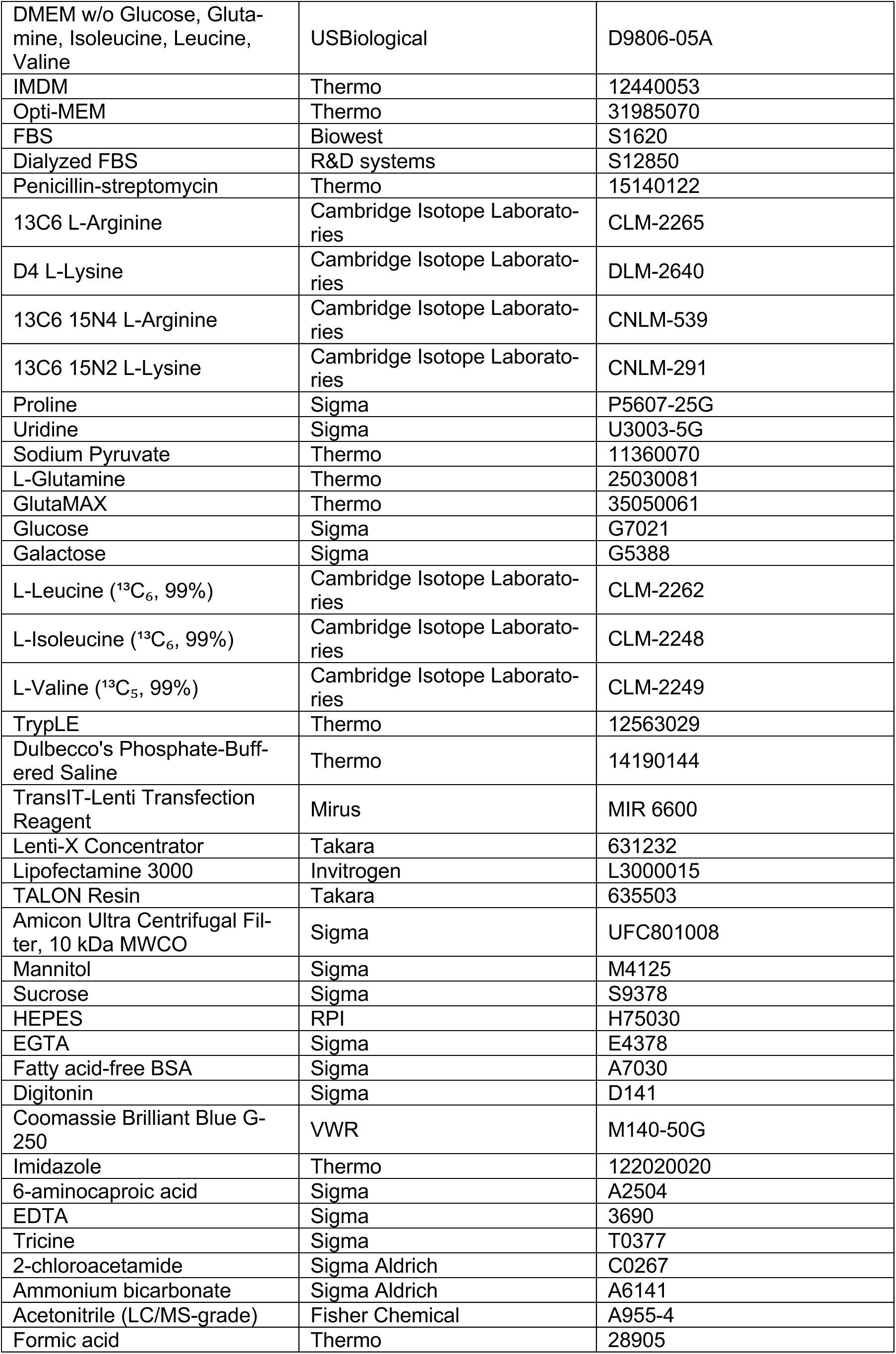
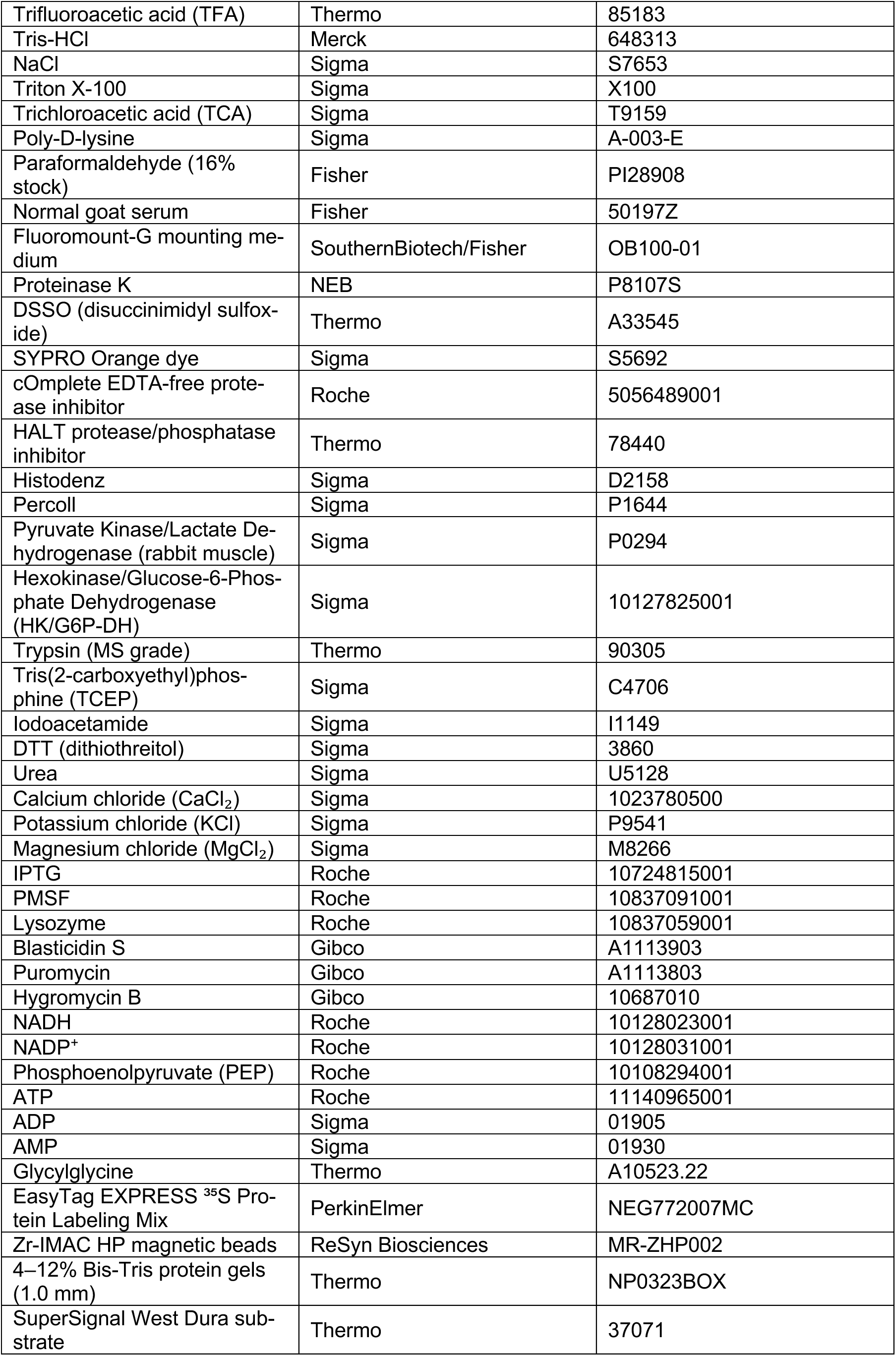
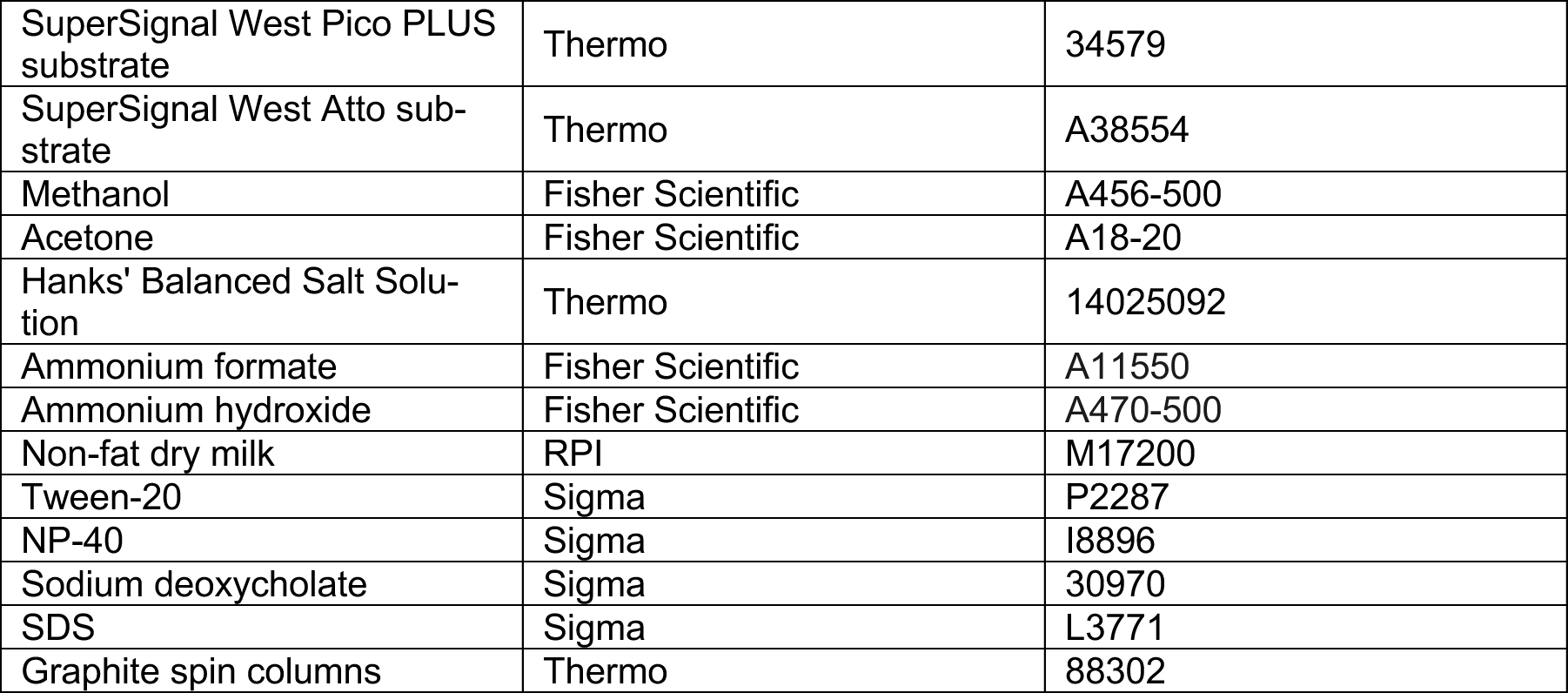

#### Critical commercial assays

| REAGENT or RESOURCE | SOURCE | IDENTIFIER |
| --- | --- | --- |
| Pierce BCA Protein Assay Kit | Thermo | 23225 |
| Peptide Clean-up Plate | Thermo | A57865 |
| Pierce Peptide Desalting Spin Columns | Thermo | 89851 |
| Quant-iT PicoGreen dsDNA Assay | Thermo | P7589 |
| High-Select TiO<sub>2</sub>/Fe-NTA SMOAC phosphopeptide enrichment kits | Thermo | A32993; A32992 |
| Q5 Site-Directed Mutagenesis Kit | New England Biolabs | E0554S |
| TnT Quick Coupled Transcription/Translation System (T7) | Promega | L2080 |

#### Deposited data

| REAGENT or RESOURCE | SOURCE | IDENTIFIER |
| --- | --- | --- |
| Phosphatase CRISPRi SILAC proteomics and phosphoproteomics (DDA) | This paper | PXD084349 |
| PGAM5-KD rescue proteomics and phosphoproteomics, MIC19 and ATP5I rescue proteomics (DIA) | This paper | PXD084307 |
| Complexome profiling study | This paper | PXD084375 |
| MIC60 AE-MS study | This paper | PXD084235 |

#### Experimental models: Cell lines

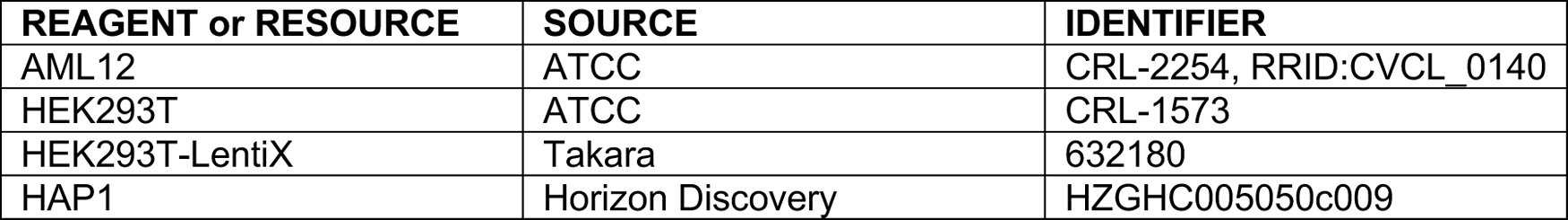

#### Oligonucleotides

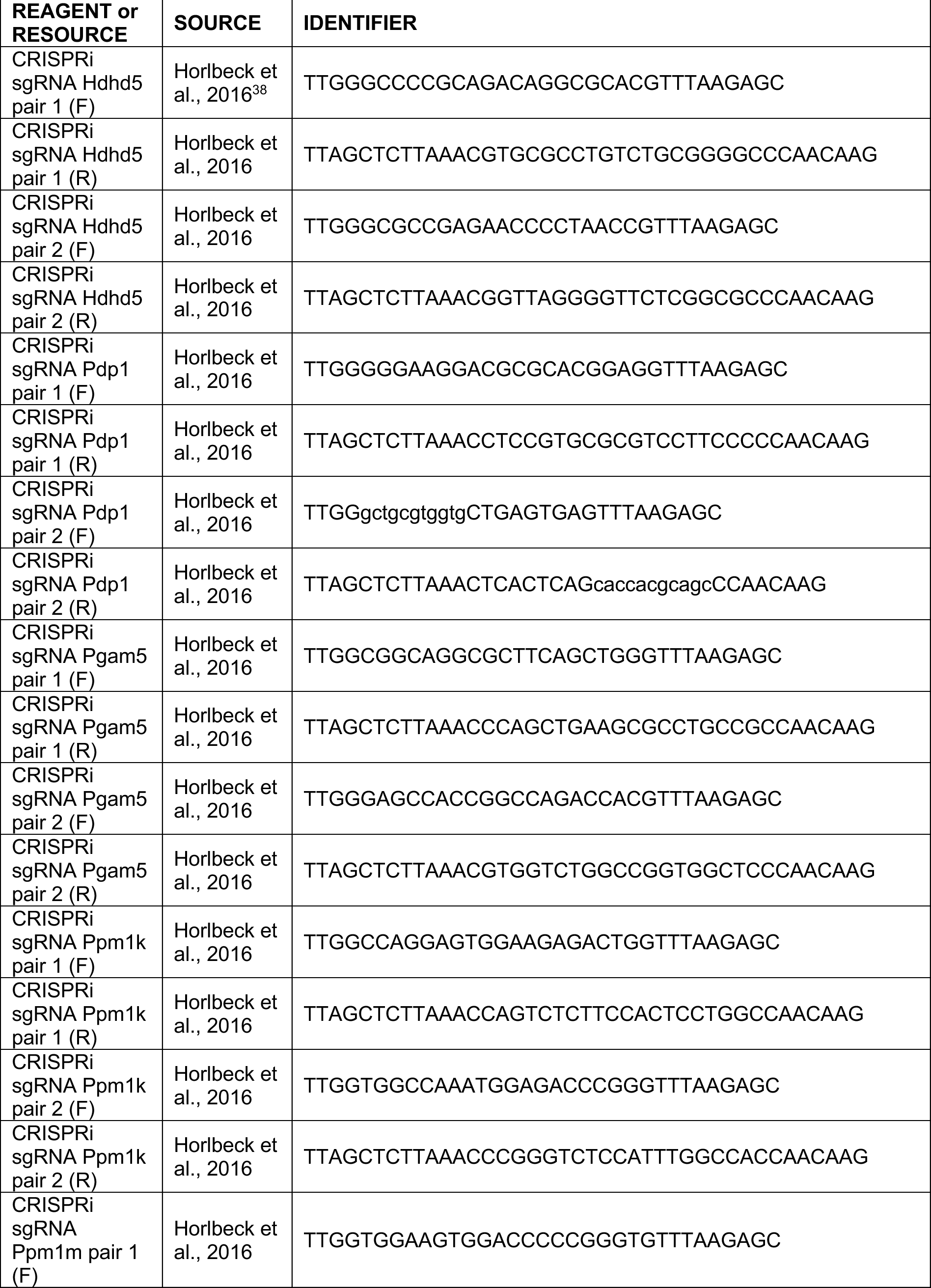
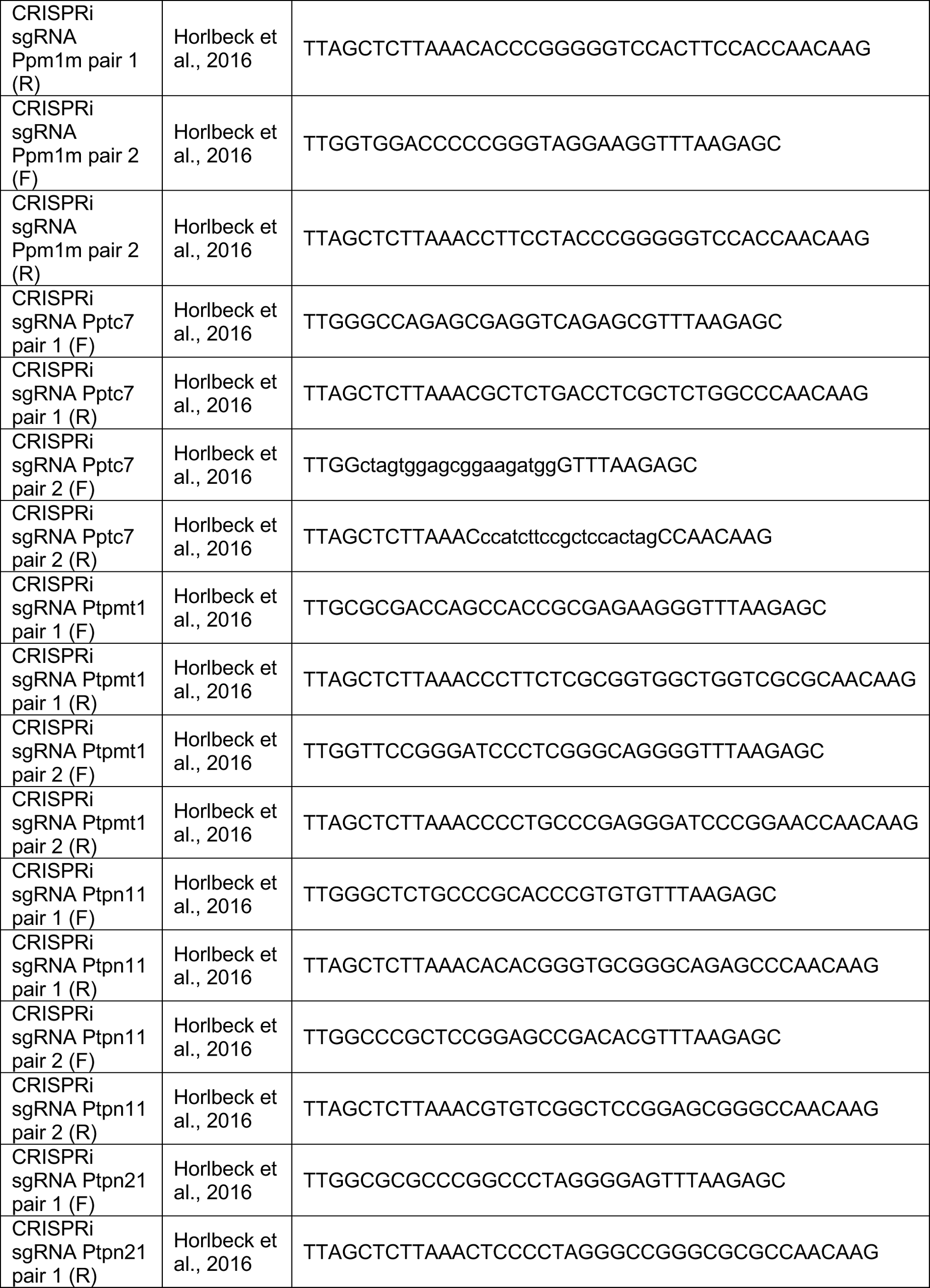
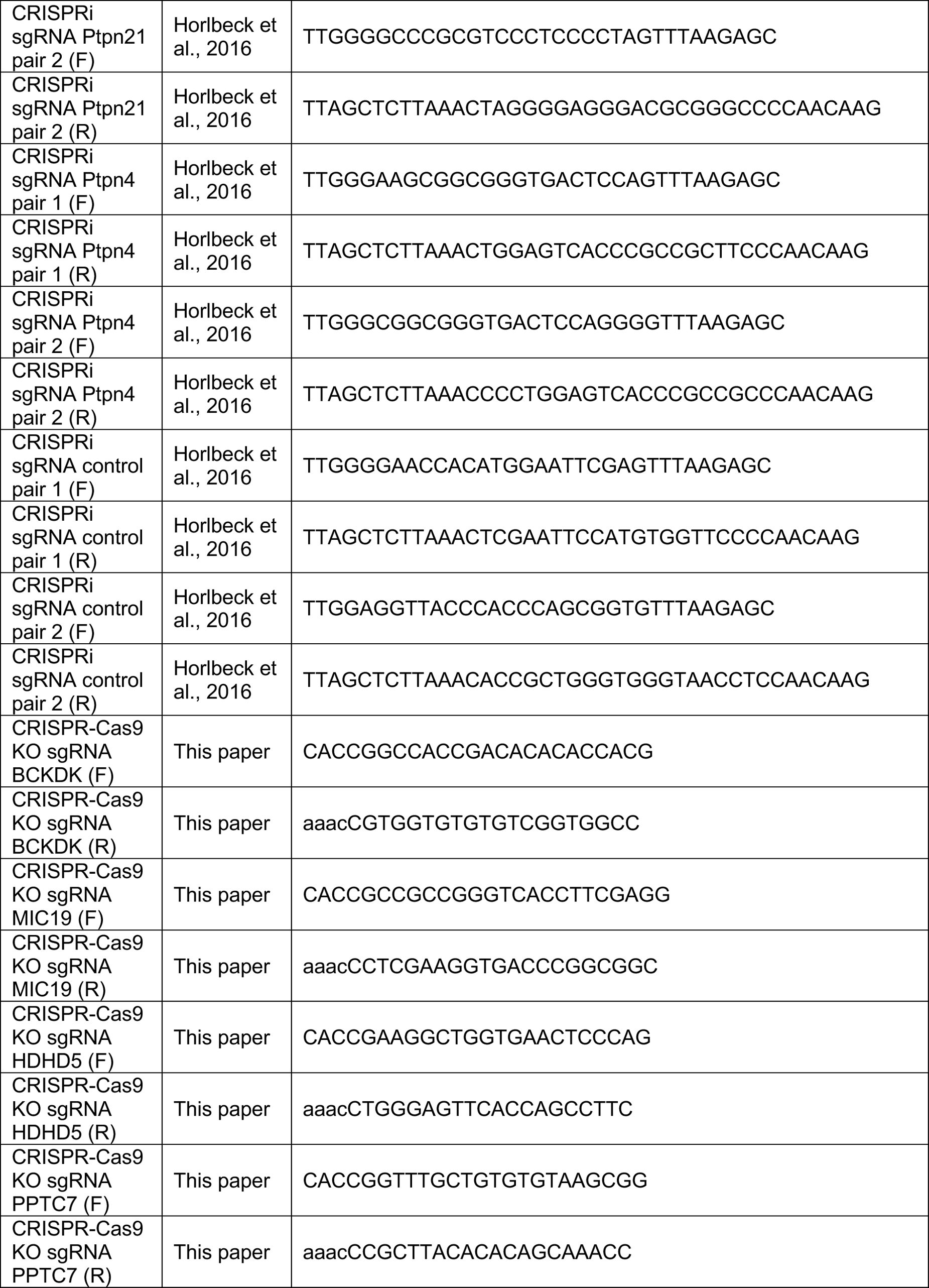

#### Recombinant DNA

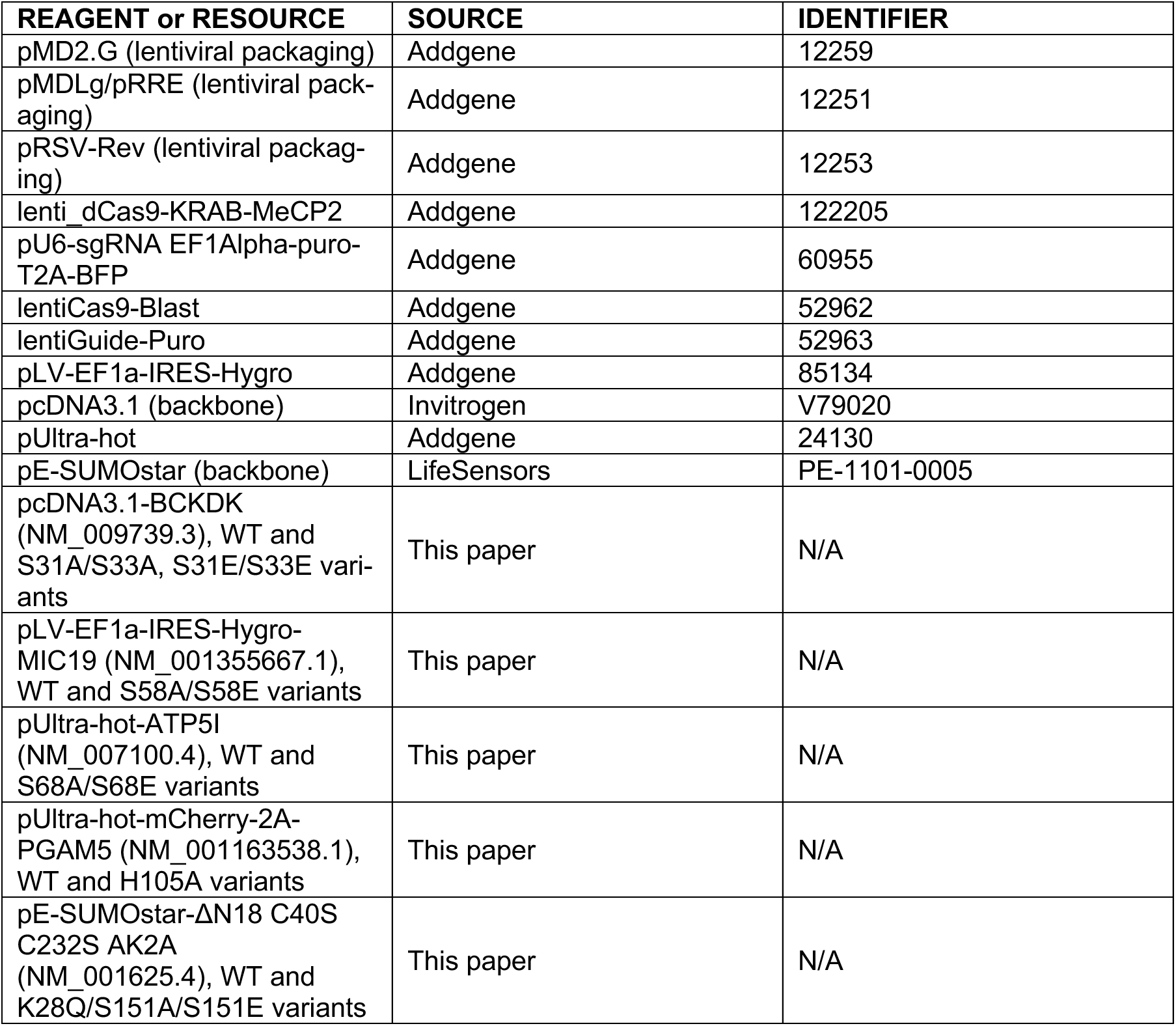

#### Software and algorithms

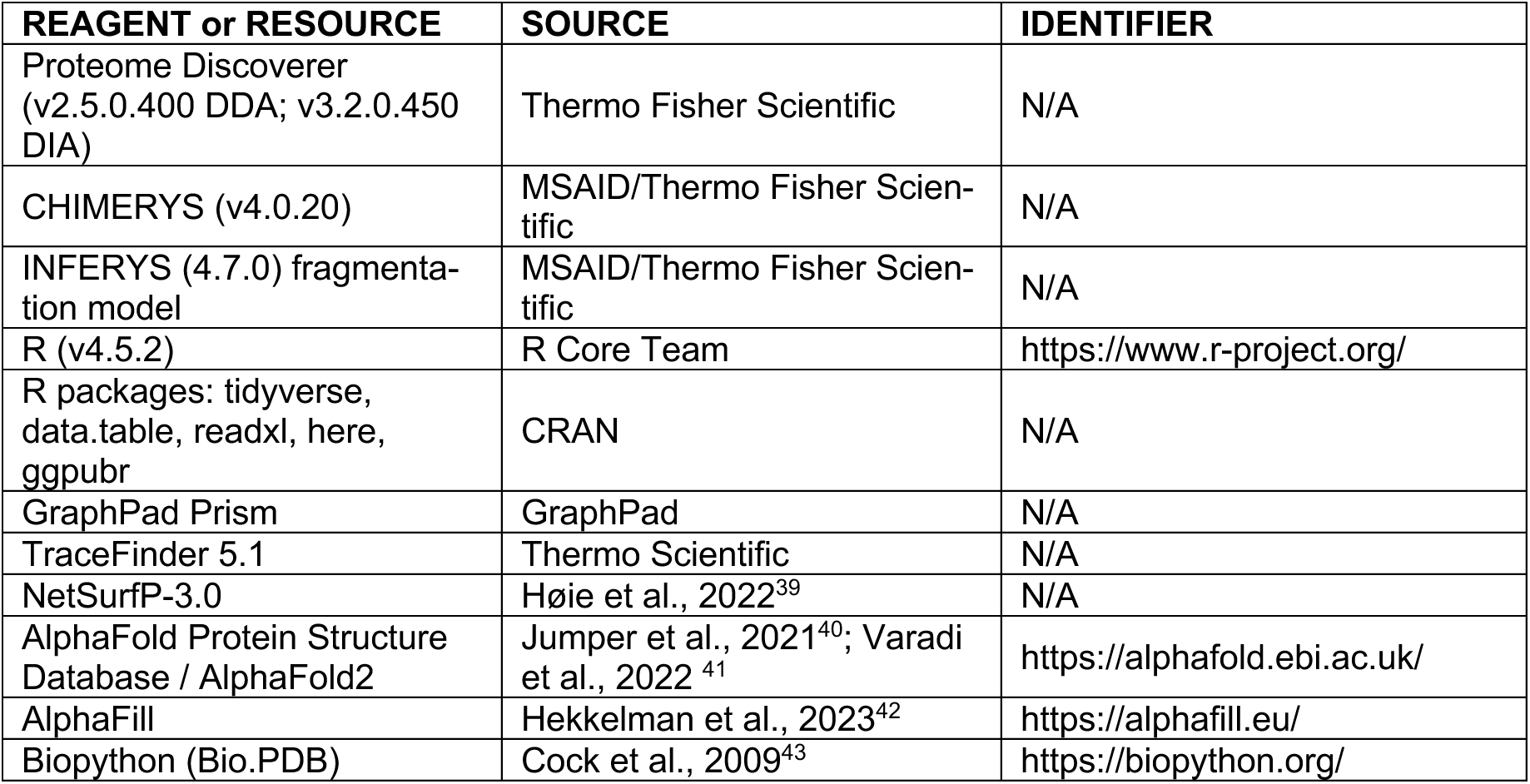
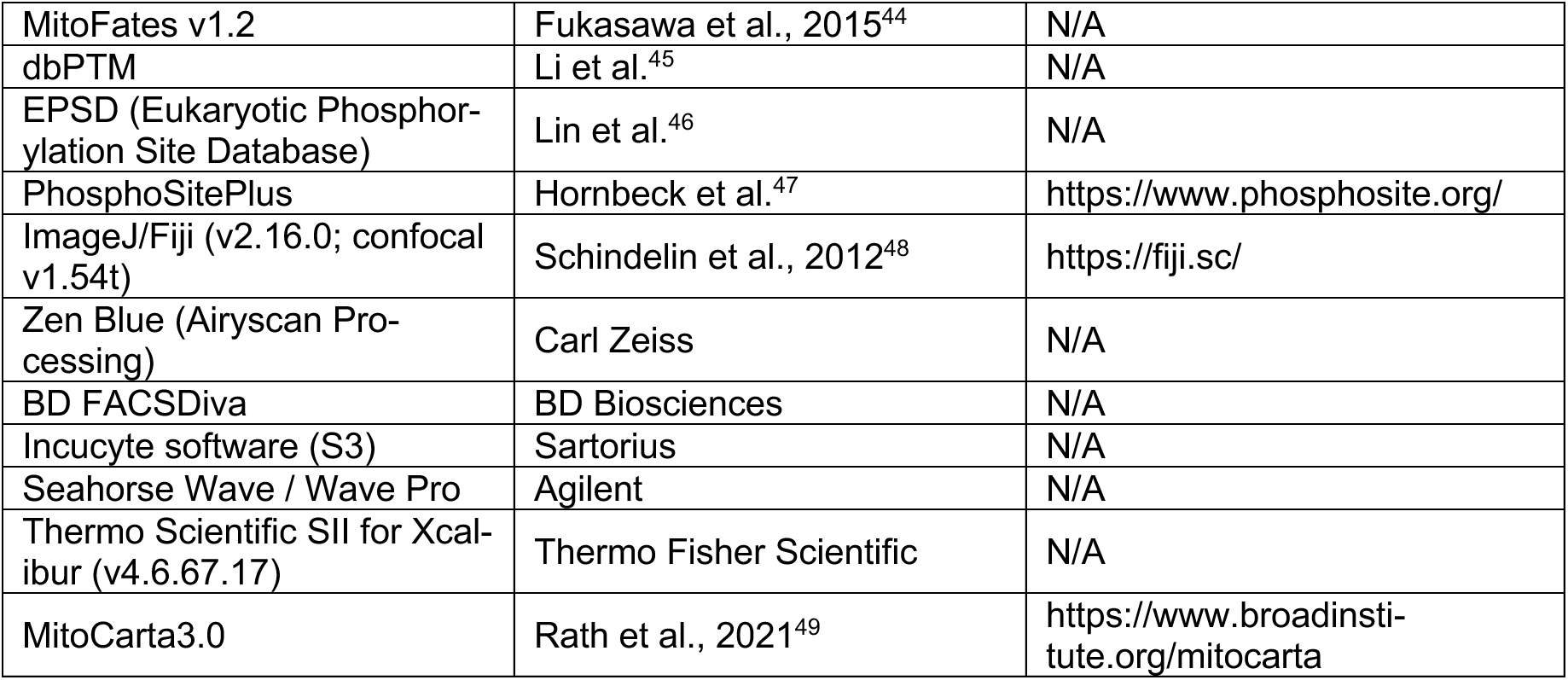

#### Other

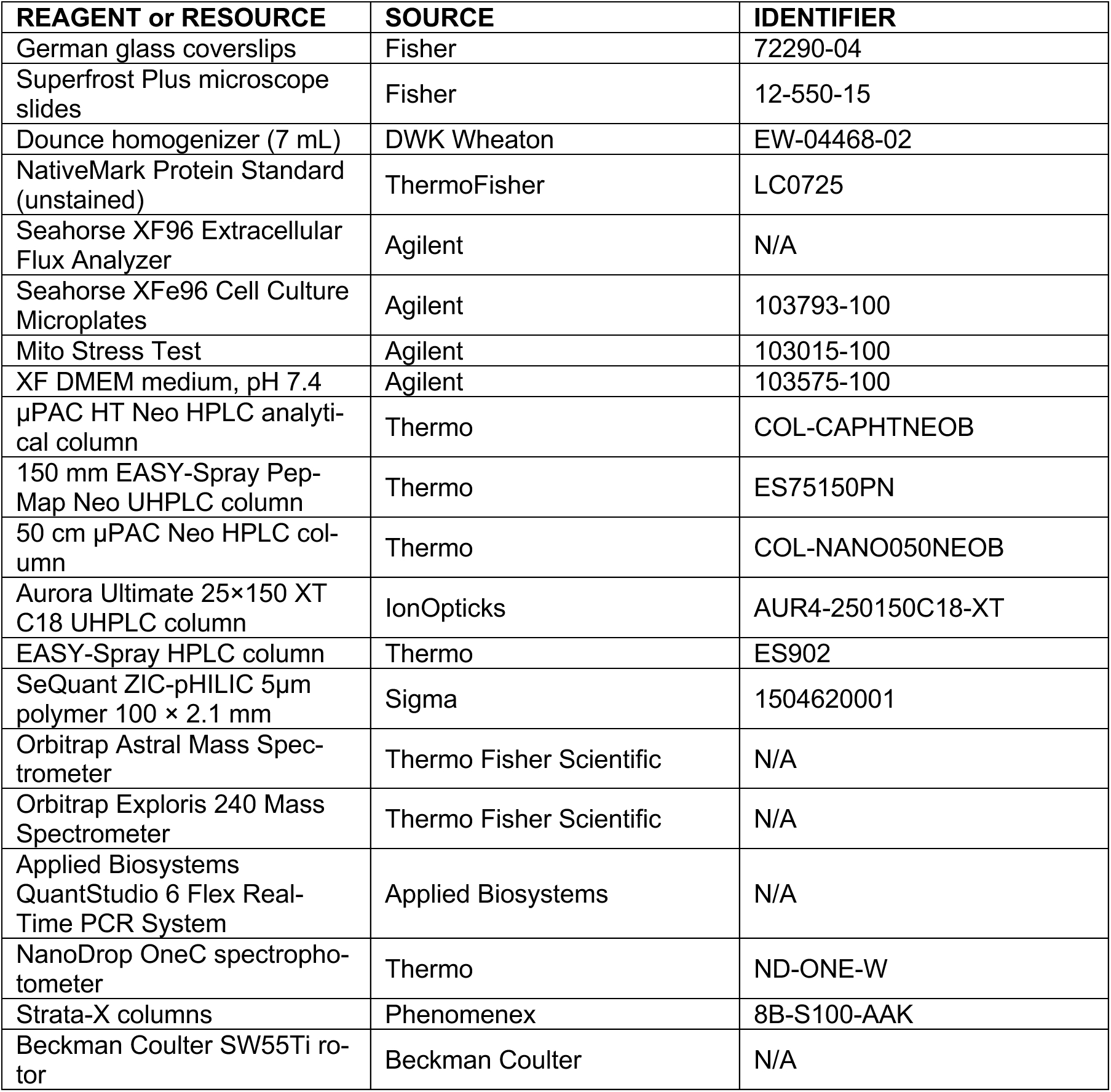
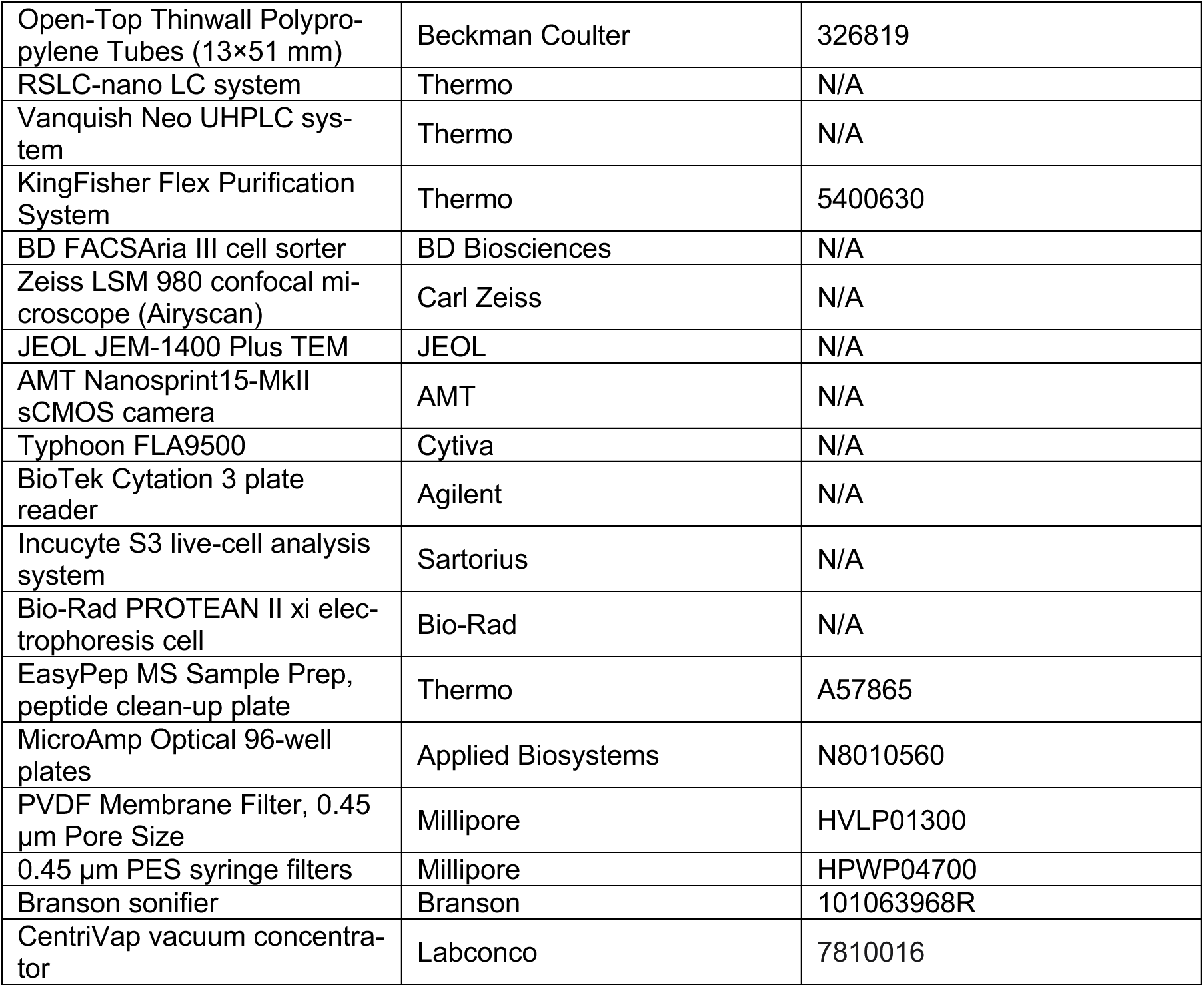

| REAGENT or RESOURCE | SOURCE | IDENTIFIER |
| --- | --- | --- |
| German glass coverslips | Fisher | 72290-04 |
| Superfrost Plus microscope slides | Fisher | 12-550-15 |
| Dounce homogenizer (7 mL) | DWK Wheaton | EW-04468-02 |
| NativeMark Protein Standard (unstained) | ThermoFisher | LC0725 |
| Seahorse XF96 Extracellular Flux Analyzer | Agilent | N/A |
| Seahorse XFe96 Cell Culture Microplates | Agilent | 103793-100 |
| Mito Stress Test | Agilent | 103015-100 |
| XF DMEM medium, pH 7.4 | Agilent | 103575-100 |
| µPAC HT Neo HPLC analytical column | Thermo | COL-CAPHTNEOB |
| 150 mm EASY-Spray Pep-Map Neo UHPLC column | Thermo | ES75150PN |
| 50 cm µPAC Neo HPLC column | Thermo | COL-NANO050NEOB |
| Aurora Ultimate 25×150 XT C18 UHPLC column | IonOpticks | AUR4-250150C18-XT |
| EASY-Spray HPLC column | Thermo | ES902 |
| SeQuant ZIC-pHILIC 5µm polymer 100 × 2.1 mm | Sigma | 1504620001 |
| Orbitrap Astral Mass Spectrometer | Thermo Fisher Scientific | N/A |
| Orbitrap Exploris 240 Mass Spectrometer | Thermo Fisher Scientific | N/A |
| Applied Biosystems QuantStudio 6 Flex Real-Time PCR System | Applied Biosystems | N/A |
| NanoDrop OneC spectrophotometer | Thermo | ND-ONE-W |
| Strata-X columns | Phenomenex | 8B-S100-AAK |
| Beckman Coulter SW55Ti rotor | Beckman Coulter | N/A |
| Open-Top Thinwall Polypropylene Tubes (13×51 mm) | Beckman Coulter | 326819 |
| RSLC-nano LC system | Thermo | N/A |
| Vanquish Neo UHPLC system | Thermo | N/A |
| KingFisher Flex Purification System | Thermo | 5400630 |
| BD FACSAria III cell sorter | BD Biosciences | N/A |
| Zeiss LSM 980 confocal microscope (Airyscan) | Carl Zeiss | N/A |
| JEOL JEM-1400 Plus TEM | JEOL | N/A |
| AMT Nanosprint15-MkII sCMOS camera | AMT | N/A |
| Typhoon FLA9500 | Cytiva | N/A |
| BioTek Cytation 3 plate reader | Agilent | N/A |
| Incucyte S3 live-cell analysis system | Sartorius | N/A |
| Bio-Rad PROTEAN II xi electrophoresis cell | Bio-Rad | N/A |
| EasyPep MS Sample Prep, peptide clean-up plate | Thermo | A57865 |
| MicroAmp Optical 96-well plates | Applied Biosystems | N8010560 |
| PVDF Membrane Filter, 0.45 µm Pore Size | Millipore | HVLP01300 |
| 0.45 µm PES syringe filters | Millipore | HPWP04700 |
| Branson sonifier | Branson | 101063968R |
| CentriVap vacuum concentrator | Labconco | 7810016 |

### RESOURCE AVAILABILITY

### Lead contact

Further information and requests for resources and reagents should be directed to and will be fulfilled by the lead contact, David J. Pagliarini.

### Materials availability

Reagents generated in this study are available from the lead contact upon request with a completed Materials Transfer Agreement.

### Data and code availability

- All mass spectrometry proteomics data (phosphoproteomics, proteomics, complexome profiling, and affinity enrichment) have been deposited to PRIDE/ProteomeXchange under the following accessions: PXD084349, PXD084307, PXD084375, PXD084235. Microscopy and all other source data reported in this paper will be shared by the lead contact upon request. Accession numbers are listed in the Key Resources Table.
- This paper does not report original code.
- Any additional information required to reanalyze the data reported in this paper is available from the lead contact upon request.

### EXPERIMENTAL MODEL AND SUBJECT DETAILS

### Cell lines and culture

AML12 mouse hepatocyte cells were maintained in DMEM/F-12 (Thermo, 11320033), HEK293T (human) cells in DMEM, high glucose (Thermo, 11965092), and HAP1 (human) cells in IMDM (Thermo, 12440053). All media were supplemented with 10% FBS (Biowest, S1620) and 1× penicillin-streptomycin (Thermo, 15140122). All cells were maintained at 37°C in a humidified incubator with 5% CO₂, and tested negative for mycoplasma.

## METHOD DETAILS

### Generation of stable cell lines

#### Lentiviral production

Lentiviral particles were generated for stable integration of either sgRNAs or gene open reading frames. HEK293T LentiX cells (Takara, 632180) were seeded in 10 cm dishes 24 h prior to transfection to achieve 80–90% confluency. Cells were co-transfected with 5 µg of the plasmid of interest along with packaging plasmids pMD2.G (Addgene, 12259), pMDLg/pRRE (Addgene, 12251), and pRSV-Rev (Addgene, 12253) at a 1:4:1 molar ratio using TransIT-Lenti Transfection Reagent (Mirus, MIR 6600) in Opti-MEM (Thermo, 31985070) according to the manufacturer’s instructions. At 48 h post-transfection, virus-containing supernatant was filtered through 0.45 µm PES filters, concentrated using Lenti-X Concentrator (Takara, 631232) at a 1:3 ratio, and incubated at 4°C with end-over-end rotation. Virus was pelleted by centrifugation at 1,500 × g for 45 min at 4°C. Pellets were resuspended in 500 µL DPBS (Thermo, 14190144), aliquoted, and stored at −80°C.

#### CRISPRi-mediated knockdown

CRISPRi knockdown was performed using a dual-component system^38^. AML12 cells stably expressing dCas9-KRAB-MeCP2 (Addgene, 122205) were generated via lentiviral transduction, blasticidin selection (2.5 µg/mL), and monoclonal isolation by limiting dilution. For sgRNA expression, annealed oligonucleotides (Key Resources Table) were cloned into the pU6-sgRNA EF1Alpha-puro-T2A-BFP plasmid (Addgene, 60955) as previously described^38^. Lentivirus was produced by combining two sgRNA plasmids 1:1 during transfection. Cells were transduced and selected with 0.4 µg/mL puromycin for 5 days, followed by recovery in antibiotic-free medium for at least 7 days before experiments.

#### CRISPR-Cas9 knockout

Monoclonal AML12 and HEK293T cells stably expressing Cas9 were generated using lentiCas9-Blast (Addgene, 52962). Knockout cell lines were established by transducing Cas9-expressing cells with lentiGuide-Puro (Addgene, 52963) containing target-specific sgRNAs (Key Resources Table). Following transduction, cells were selected with 0.5 µg/mL puromycin for 5 days and recovered in antibiotic-free medium before monoclonal isolation.

#### Stable rescue and overexpression cell lines

Gene rescue and overexpression cell lines were generated using pcDNA3.1 (BCKDK, transient transfection), pUltra-hot (Addgene, 24130; PGAM5, ATP5I, see FACS), or pLV-EF1a-IRES-Hygro (Addgene, 85134; MIC19) vectors containing the gene of interest (Key Resources Table). Cells transduced with pLV-EF1a-IRES-Hygro lentivirus were selected with hygromycin (300 µg/mL).The following cDNA sequences were used: BCKDK (NM_009739.3), MIC19 (NM_001355667.1), ATP5I (NM_007100.4), and PGAM5 (NM_001163538.1), or point-mutant variants thereof.

#### Fluorescence-activated cell sorting

Cells transduced with pUltra-hot vectors encoding PGAM5 or ATP5I, or a variant thereof, were sorted on a BD FACSAria III cell sorter (BD Biosciences) equipped with an 85 µm nozzle. Cells were sorted for mCherry positivity using a 600 nm longpass dichroic mirror into a 610/20 nm bandpass filter.

#### Isolation of crude mitochondria

Unless otherwise stated, mitochondrial isolates were obtained from cultured cells by differential centrifugation to preserve organelle integrity. Several days before isolation, cells were plated and grown to approximately 90% confluency. On the day of harvest, culture medium was removed and cells were washed once with DPBS, then detached using a cell scraper in fresh ice-cold PBS and transferred to prechilled 50 mL conical tubes on ice. After collecting all plates of the same sample group, the cell suspension was centrifuged at 600 × g for 10 min at 4°C, washed once with 10 mL ice-cold PBS, and pelleted again at 600 × g for 5 min. The pellet was resuspended in 3 mL ice-cold mitochondrial isolation buffer (210 mM mannitol, 70 mM sucrose, 5 mM HEPES pH 7.4, 1 mM EGTA) freshly supplemented with protease inhibitors (Roche, cOmplete EDTA-free, 05056489001). A separate aliquot of the same buffer containing 0.5% fatty acid–free BSA was used for resuspension and homogenization. Cells were transferred to a prechilled 7 mL Dounce homogenizer (DWK Wheaton, EW-04468-02) and disrupted on ice with 30–35 strokes of a tight pestle. The homogenate was centrifuged at 600 × g for 10 min at 4°C to remove nuclei and debris. The supernatant was transferred to fresh tubes and spun at 7,000 × g for 10 min at 4°C. The resulting pellet was the mitochondria-enriched fraction and the supernatant the cytosolic fraction. The mitochondrial pellet was washed once with 200 µL prechilled isolation buffer and centrifuged again at 7,000 × g for 10 min at 4°C. The final pellet was gently resuspended in 200 µL isolation buffer using wide-bore tips. Protein content was determined using a Pierce BCA Protein Assay Kit (Thermo, 23225) and mitochondria were aliquoted. The crude mitochondrial fraction was used immediately or flash-frozen and stored at −80°C. If flash-frozen, aliquoted mitochondria were centrifuged again at 7,000 × g for 10 min at 4°C and all supernatant removed.

### Proteomics and phosphoproteomics

#### SILAC-labeled phosphatase perturbation

Prior to lentiviral transduction of CRISPRi sgRNAs, AML12 dCas9-KRAB-MeCP2 monoclonal cells were cultured in SILAC Advanced DMEM/F-12 Flex Media (5% dialyzed FBS (R&D Systems, S12850), 100 mg/L L-proline, 50 µg/mL uridine, 25 mM glucose, 1 mM pyruvate, 2 mM L-glutamine) supplemented with either medium (¹³C₆ L-arginine (CIL, CLM-2265); D4 L-lysine (CIL, DLM-2640)) or heavy (¹³C₆ ¹⁵N₄ L-arginine (CIL, CNLM-539); ¹³C₆ ¹⁵N₂ L-lysine (CIL, CNLM-291)) isotopes, for a minimum of five passages, and confirmed to have near-complete SILAC incorporation by mass spectrometry. Following transduction with either control, non-targeting sgRNAs (medium) or phosphatase-targeted sgRNAs (heavy) for 48 h, cells were selected with puromycin (0.4 µg/mL) for 7 days and expanded for a further 7 days. For harvesting, cells were counted and combined 1:1 (medium:heavy) before crude mitochondrial isolation using nitrogen cavitation for 10 min as described previously^50^, continuing as described above; samples were flash-frozen and stored at −80°C.

#### Sample preparation

Crude mitochondrial pellets, or whole cell lysates, where indicated, were thawed on ice and resuspended in 250 µL lysis buffer (8 M urea, 100 mM Tris-HCl pH 8.0, 1× HALT protease/phosphatase inhibitor (Thermo, 78440)). Samples were sonicated with a probe sonicator (2 × 10 s pulse, 10% output) and clarified by centrifugation at 16,500 × g for 15 min at 4°C. Protein concentrations were determined by BCA assay (Thermo, 23225). Samples were reduced with tris(2-carboxyethyl)phosphine (10 mM final) and alkylated with 2-chloroacetamide (40 mM final) for 1 h at room temperature (1,000 rpm). Proteins were digested with trypsin (Thermo, 90305) at a 1:25 enzyme-to-protein ratio, ensuring a final urea concentration below 2 M. CaCl₂ was added to 1 µM final and the volume adjusted with 100 mM Tris-HCl pH 8.0. Digestion proceeded overnight (16–20 h) at 37°C and 1,000 rpm.

#### Peptide desalting

For analysis on the Orbitrap Exploris 240, Strata-X columns (Phenomenex, 8B-S100-AAK) were prepared on the day of use. Columns were conditioned with 2 column volumes of 100% ACN followed by 2 column volumes of 0.1% TFA. Samples were acidified with TFA to pH < 3.0 and centrifuged at 16,000 × g for 5 min. Supernatants were loaded, washed twice with 2 column volumes of 0.1% TFA, and eluted with 80% ACN/0.1% TFA into low-protein-binding tubes. Five percent of the eluate was retained for proteomics analysis and the remainder for phosphopeptide enrichment. For analysis on the Orbitrap Astral, peptides were desalted using Pierce Peptide Desalting Spin Columns (Thermo, 89851) per the manufacturer’s instructions with minor modifications (activated twice with 100% ACN, equilibrated twice with 0.1% TFA at 5,000 × g for 1 min; loaded and washed at 3,000 × g; eluted twice with 50% ACN/0.1% TFA). In both workflows, eluates were dried in a SpeedVac and resuspended in 0.2% formic acid, or stored at −80°C.

#### Phosphopeptide enrichment

Phosphopeptide enrichment of Orbitrap Exploris 240 samples used the High-Select Sequential Metal Oxide Affinity Chromatography (SMOAC) method per the manufacturer’s instructions (Thermo, A32993 and A32992), with 500 µg peptides as input; enriched peptides were cleaned up on graphite spin columns (Thermo, 88302). Phosphopeptide enrichment of Orbitrap Astral samples used Zr-IMAC (ReSyn Biosciences, MR-ZHP002) with 100 µg peptides as input, at a 3.3:1 bead-to-peptide ratio as previously described^51^, on a King-Fisher Flex Purification System (Thermo, 5400630). Peptides were resuspended in 0.2% formic acid prior to analysis.

#### Peptide quantification and injection

Peptide concentrations were quantified after resuspension in 0.2% formic acid using a NanoDrop OneC spectrophotometer (Thermo) with the Protein A205 method. For LC–MS acquisition, 1 µg peptide was injected for the Exploris 240 and 200 ng for the Orbitrap Astral.

#### Liquid chromatography

For the Orbitrap Exploris 240, peptides were separated on an RSLC-nano system (Thermo) using 120 min methods on an EASY-Spray column (Thermo, ES902; 75 µm × 250 mm) with buffer A (0.1% FA in water) and buffer B (80:20 ACN:water, 0.1% FA) at 0.3 µL/min (maximum 800 bar). The gradient differed by sample type. Phosphoproteomics: 4% B at 3 min, 8% B at 5 min, 40% B at 100 min, ramp to 99% B over 105–110 min (column wash), and return to 4% B over 111–120 min (re-equilibration). Proteomics: 4% B at 3 min, 20% B at 60 min, 45% B at 100 min, ramp to 99% B over 105–110 min, and return to 4% B by 111 min (held through the 120 min method). Controlled flow-ramp limits (NC pump ±0.3 µL/min²; loading pump ±5 µL/min²) and two automated trap-wash cycles were applied; methods ran under Thermo Scientific SII for Xcalibur (v4.6.67.17). For the Orbitrap Astral, peptides were separated on a Vanquish Neo UHPLC system (Thermo) with buffers A and B, using a distinct method per sample type. (i) PGAM5-rescue proteomics (24 min gradient): 200 ng was loaded at 20 µL/min onto a trap column (300 µm × 0.5 cm, backward-flush mode) and eluted onto a 150 mm EASY-Spray PepMap Neo column (Thermo, ES75150PN) at 0.5 µL/min; 4% B to 0.5 min, 8% B at 0.6 min, 22.5% B at 13.9 min, 35% B at 20.8 min, 55% B at 21.2 min, 99% B wash from 21.2–23.9 min, 87.5% B at 24 min, then equilibration. (ii) PGAM5-rescue phosphoproteomics (45 min gradient) on a 50 cm µPAC Neo column (Thermo, COL-NANO050NEOB): 1% B to 0.1 min, 2% B to 0.3 min, 8% B at 3.2 min, 22.5% B at 28.2 min, 45% B at 36.1 min, then 99% B wash from 37.6–45 min. (iii) MIC19 and ATP5I proteomics on an Aurora Ultimate 25 × 150 XT C18 column (IonOpticks, AUR4-250150C18-XT) at 40°C in trap-and-elute mode (NanoCap regime, 0.5 µL/min, 1,500 bar): 3% B, 8% B at 1.1 min, 11.4% B at 2.9 min, 21% B at 12.1 min, 43.8% B at 30.7 min, 99% B wash from 31.7–36.0 min, then re-equilibration (fast equilibration 0.8 µL/min, 400 bar; two 800-bar trap washes).

#### Mass spectrometry data acquisition

On the Orbitrap Exploris 240, data were acquired in DDA mode (120 min; positive polarity, spray voltage +1,800 V, ion transfer tube 275°C; EASY-IC). Phosphoproteomic MS1: Orbitrap 120,000 resolution, m/z 375–1,500, RF lens 70%, AGC 300%, max IT 100 ms; ddMS2 in cycle-time mode (3 s), isolation 1.6 m/z, HCD 25%, Orbitrap 30,000, AGC 25%. Proteomic MS1 differed in RF lens 80%, max IT 50 ms, intensity threshold 1.0 × 10⁵, dynamic exclusion 60 s, and MS2 AGC 75%. On the Orbitrap Astral, data were acquired in DIA mode (positive polarity, +1,500 V, ion transfer tube 280°C; no FAIMS; EASY-IC). Proteomics: MS1 Orbitrap 240,000 (m/z 380–980), RF lens 40%, AGC 5.0 × 10⁶, max IT 3 ms; DIA in Astral mode with 2 m/z windows (300 events), HCD 25%, AGC 5.0 × 10⁴, max IT 3 ms, 0.6 s cycle. Phosphoproteomic DIA modified MS1 AGC to 2.0 × 10⁶ and MS2 to 4 m/z windows, RF lens 45%, max IT 20 ms.

#### Recombinant protein expression and purification

A previously described^52^ codon-optimized ΔN18 C40S C232S AK2A (NM_001625.4) was cloned into the pE-SUMOstar vector with a 5′ SUMO-tag coding sequence. Point mutations were generated using the Q5 Site-Directed Mutagenesis Kit (New England Biolabs) and plasmids were transformed into Rosetta(DE3)pLysS cells (Sigma, 70956-3). Cells were induced at OD600 0.6 with 0.1 mM IPTG for 18 h at 16°C before pelleting (4,000 × g, 20 min, 4°C). Cells were resuspended in lysis buffer (50 mM HEPES pH 7.4, 500 mM NaCl, 5% glycerol, 1 mM PMSF, 1 mM DTT, 1 mg/mL lysozyme) and sonicated on ice with a Branson sonifier (75% amplitude, 20 s on/1 min rest, two cycles). Lysates were clarified (15,000 × g, 30 min, 4°C). The soluble fraction was incubated with 3 mL TALON metal-affinity resin (Takara, 635503) for 2 h at 4°C. The resin was washed with buffer containing increasing imidazole (10, 20, 30 mM) and eluted with 25 mL buffer containing 100 mM imidazole. Buffer exchange and concentration used a 10 kDa MWCO filter (Sigma, UFC801008) into exchange buffer (50 mM HEPES pH 7.4, 500 mM NaCl, 5% glycerol, 1 mM DTT). Purity was verified by SDS-PAGE. Purified proteins were stored fresh (4°C) or in 50% glycerol at −80°C.

#### In vitro enzyme activity assays

AK2 activity with AMP as the variable substrate was assayed by coupling to pyruvate kinase (PK) and lactate dehydrogenase (LDH; Sigma, P0294): ADP produced by AK2 drives PK conversion of phosphoenolpyruvate to pyruvate, which LDH uses to oxidize NADH, decreasing absorbance at 340 nm (50 mM Tris-HCl pH 7.5, 50 mM KCl, 4 mM MgCl₂, 0.006% BSA, 0.25 mM NADH, 1 mM PEP, 1 mM ATP). AK2 activity with ADP as the variable substrate was coupled to hexokinase (HK) and glucose-6-phosphate dehydrogenase (G6PDH; Sigma, 10127825001): ATP generated by AK2 supports glucose phosphorylation, reducing NADP⁺ to NADPH and increasing absorbance at 340 nm (58 mM glycylglycine pH 7.4, 10 mM MgCl₂, 0.006% BSA, 0.25 mM NADP⁺, 20 mM glucose). Assays were performed at 25°C in a BioTek Cytation 3 plate reader (200 µL reactions, 1 µg enzyme per well, technical duplicate with no-enzyme and no-coupling-mix controls). Two independent protein preparations were assayed, with ≥2 replicates of forward and reverse assays each. Initial velocity was the slope of A340 over the first 60 s. Velocities (ΔA340 min⁻¹, magnitude of NAD(P)H change) were fit to v = Vmax·[S]/(Km + [S]) by non-linear least-squares in GraphPad Prism (Km, Vmax constrained positive). K28Q showed no detectable activity; for S151E, weakened affinity left Km poorly constrained. Velocities were converted to molar NAD(P)H turnover using ε = 6,220 M⁻¹cm⁻¹ and the 0.625 cm path length.

#### Differential scanning fluorimetry

Protein melting curves were performed essentially as described^53^. 20 µL reactions contained 4 µM purified protein and 5× SYPRO Orange (Sigma, S5692) in 50 mM HEPES pH 7.4, 150 mM NaCl, in MicroAmp Optical 96-well plates. Fluorescence was measured in the ROX channel during a 25→95°C ramp at 0.015°C/s on an Applied Biosystems QuantStudio 6 Flex. Tm values were calculated by fitting the transition to a Boltzmann signmoidal fit model.

#### Immunofluorescence and confocal imaging

Cells were seeded at 2.5 × 10⁴ per well in 24-well plates on coverslips. For AML12 cells transiently transfected with BCKDK constructs, cells were transfected at ∼60% confluency with Lipofectamine 3000 (Invitrogen, L3000015; 0.5 µg DNA) in Opti-MEM and fixed 24 h later with 4% PFA (Fisher, PI28908) for 5 min at 37°C. AML12 MIC19-rescue cells were grown overnight prior to 4% PFA fixation. Cells were washed twice with PBS, permeabilized and blocked in 5% normal goat serum in PBS-T (0.03% Triton X-100) for ≥30 min. Cells were incubated with primary antibodies against FLAG (CST, 14793; 1:1000) and HSP60 (EnCor, CPCA-HSP60; 1:500) overnight at 4°C, then with Alexa Fluor 488 goat anti-rabbit IgG (Invitrogen, A11008; 1:500) and Alexa Fluor 647 goat anti-chicken IgY (Invitrogen, A32933; 1:500) for 1 h. Coverslips were mounted on slides (Fisher, 12-550-15) using Fluoromount-G.

Images were acquired on a Zeiss LSM 980 confocal microscope with an Airyscan detector using a Plan-Apochromat 63×/1.40 NA objective. MIC19 samples were imaged in Airyscan Super-Resolution mode (pinhole ∼5 AU, 8× zoom, 4× line averaging; 42.5 nm XY; 20 z-sections at 0.15 µm). BCKDK samples used standard confocal (pinhole 1.18 AU, 2× zoom, 4× frame averaging; 65.8 nm XY; 18 z-sections at 0.27 µm). Airyscan data were processed in Zen Blue; subsequent processing used ImageJ (v1.54t).

#### [35S] mitochondrial import assay

Import assays were performed as previously described^54^. WT BCKDK and variants cloned into pcDNA3.1 with an upstream T7 promoter were used for in vitro transcription/translation with EasyTag EXPRESS ³⁵S Protein Labeling Mix (PerkinElmer, NEG772007MC) and the TnT Quick Coupled Transcription/Translation System (T7; Promega, L2080). Crude mitochondria from HEK293T cells (20 µg per assay) were used; import proceeded for 1, 3, and 10 min at 30°C. Samples were placed on ice, centrifuged (8,000 × g, 5 min, 4°C), and resuspended in isolation buffer containing proteinase K, or proteinase K + 0.5% Triton X-100 for the 10 min sample, for 20 min on ice. The digest was inhibited with 1 mM PMSF before pelleting (10,000 × g, 5 min). Pellets were resuspended in LDS buffer + 50 mM DTT, boiled, and analyzed by SDS-PAGE. Gels were exposed to an autoradiography screen for 24 h and developed on a Typhoon imager (PMT 800 V, 100 µm resolution, Typhoon FLA9500, Cytiva).

#### Protein electrophoresis and western blotting

Cell pellets were lysed in RIPA buffer (50 mM Tris-HCl pH 7.4, 150 mM NaCl, 1% NP-40, 1 mM EDTA, 1 mM EGTA, 0.1% SDS, 0.5% sodium deoxycholate, protease/phosphatase inhibitors) on ice for 45 min, clarified (16,500 × g, 20 min, 4°C), and quantified by BCA. Samples were prepared in LDS buffer + 50 mM DTT, heated at 95°C for 5 min; 10 µg was loaded per lane of 4–12% Bis-Tris gels (Thermo, NP0323BOX) and resolved at 200 V for 35 min in 1× MES buffer. Proteins were transferred to methanol-activated PVDF at 20 V for 1 h. For BN-PAGE immunoblots, gels were incubated in denaturing buffer (300 mM Tris pH 8.6, 100 mM acetic acid, 1% SDS) for 20 min, transferred to PVDF overnight at 4°C in 50 mM tricine/7.5 mM imidazole, and destained with methanol. Membranes were blocked in 5% milk/TBST and probed with primary antibodies (listed in the Key Resources Table) overnight at 4°C, then HRP-conjugated secondaries (CST 7074/7076; 1:2000) for 1 h. Signals were detected with SuperSignal West Dura, Pico PLUS, or Atto (Thermo, 37071/34579/A38554).

### Branched-chain amino acid pulse-chase measurement

#### In vitro pulse-chase and sample preparation

Monoclonal BCKDK-knockout HEK293T cells were seeded in 10 cm dishes and transfected with rescue pcDNA3.1 constructs (15 µg) using Lipofectamine 3000. At 24 h, cells were seeded in 6-well plates in complete DMEM. PPTC7- and HDHD5-knockout cells were similarly seeded. The next morning, medium was swapped to heavy-BCAA DMEM (USBiological, D9806-05A; 10% dialyzed FBS, 25 mM glucose, 2 mM L-glutamine, 1× penicillin/streptomycin, 150 µM each of ¹³C₆ L-leucine (CLM-2262), ¹³C₆ L-isoleucine (CLM-2248), ¹³C₅ L-valine (CLM-2249)) for 3 h, then to DMEM without BCAAs. Samples were harvested at time 0 and at chase timepoints, normalized by cell count, extracted in 500 µL ice-cold 80% MeOH, vortexed, and held at −80°C ≥15 min. After centrifugation (21,000 × g, 10 min, 4°C), 400 µL supernatant was dried, resuspended in 50% ACN, clarified, and protein-quantified by BCA.

#### LC-MS data acquisition

Amino acids were separated by HILIC (SeQuant ZIC-pHILIC, 5 µm, 100 × 2.1 mm; Sigma, 1504620001) on a Vanquish UHPLC (0.3 mL/min, 35°C). Mobile phase A was ACN/water (1:1) with 10 mM ammonium formate/0.1% FA; B was ACN/water (9:1) with 10 mM ammonium formate/0.1% FA. The gradient held 100% B (2 min), decreased to 70% B (10 min) and 40% B (15 min), held to 17 min, and returned to 100% B by 20 min (25 min run). Detection used an Orbitrap Exploris 240 in PRM mode with H-ESI, alternating positive (3,500 V) and negative (2,500 V) polarity, triggering targeted product-ion scans of 22 amino acids plus three heavy internal standards (L-isoleucine, L-leucine, L-valine) within defined RT windows (0.4 m/z isolation; HCD 50/100/150%; 30,000 resolution; centroid).

#### Transmission electron microscopy

AML12 and HAP1 cells were seeded on coverslips (Electron Microscopy Sciences, 72290-04). Cells were washed with warm 0.15 M cacodylate buffer and fixed (2.5% glutaraldehyde, 2% paraformaldehyde, 0.15 M cacodylate pH 7.4, 2 mM CaCl₂) for 10 min at 37°C, then overnight at room temperature. Coverslips were processed on a Leica EM TP: rinsed in 1× HBSS, secondary-fixed in 1% osmium tetroxide/HBSS (1 h), rinsed, and stained in 2% aqueous uranyl acetate (8 h, 4°C). Samples were dehydrated in graded acetone and infiltrated into Epon 812 (Electron Microscopy Sciences), cured at 60°C for 72 h. Coverslips were dissolved in 42% hydrofluoric acid and cells mounted for en face sectioning. Thin sections (70 nm) were post-stained with uranyl acetate and Reynolds lead citrate and imaged on a JEOL JEM-1400 Plus TEM at 120 kV with an AMT Nanosprint15-MkII sCMOS camera. Mitochondria were binned based on morphological characterization as performed previously^19^. Cristae density was quantified by blinded manual tracing in Fiji (ImageJ v2.16.0) as described^55^.

#### Isolation of pure mitochondria and cytosolic fractions

For pure mitochondrial fractions, crude mitochondria were prepared as above and layered on a Histodenz/Percoll step gradient (35% Histodenz, 17% Histodenz, 6% Percoll) in Thinwall Polypropylene Tubes (Beckman Coulter, 326819). Gradients were centrifuged in an SW55Ti rotor at 19,000 rpm for 45 min at 4°C. Purified mitochondria were collected from the 17%/35% interface, diluted in 30 mL isolation buffer, re-pelleted, and resuspended for quantification by BCA, aliquoted, snap-frozen, and stored at −80°C. Cytosolic fractions were prepared by three rounds of 21,000 × g clarification followed by TCA precipitation (¼ volume 100% TCA, −20°C overnight; 21,000 × g, 15 min; 100% acetone wash; air-dried; stored −20°C).

#### Proteinase K protection assay

A proteinase K protection assay assessed the submitochondrial localization of PGAM5. Crude mitochondria from AML12 cells were resuspended in isolation buffer (no BSA or protease inhibitors) at ∼50 µg/µL, quantified by BCA, and diluted to 2 µg/µL. 2× assay solutions were prepared on ice with proteinase K (NEB, P8107S; 0.8 U/µL) and 10% digitonin: buffer alone, PK alone, and PK with 0.03%, 0.1%, 0.3%, or 1% digitonin. Mitochondria (40 µg) were incubated with 1× assay buffer on ice for 30 min, followed by 6.5 mM PMSF and 25% TCA. Samples were incubated on ice for 1 h, pelleted (16,500 × g, 10 min, 4°C), acetone-washed, air-dried, resuspended in LDS/50 mM DTT, and separated by SDS-PAGE. Blots were probed for TOM20, TOM70, PGAM5, TIMM50, ATP5A, and HSP60 (Key Resources Table).

### Complexome profiling

#### Sample preparation

Crude mitochondria were resuspended in solubilization buffer (50 mM imidazole, 500 mM 6-aminocaproic acid, 1 mM EDTA) at 10 µL per 100 µg protein. Digitonin (Sigma, D141-500MG) was added to a 6 g/g detergent-to-protein ratio, incubated on ice for 20 min, and cleared (20,000 × g, 20 min, 4°C). The supernatant was quantified by BCA and supplemented with glycerol (5% final) and Coomassie G-250 (VWR, M140-50G) at an 8 g/g detergent-to-dye ratio.

#### Blue native PAGE

Complexes were separated on a Bio-Rad PROTEAN II xi system on self-cast 4–13% native gels^56^. Wells were rinsed with BN anode buffer (25 mM imidazole pH 7.0) before loading 50 µg lysate and 10 µL NativeMark ladder (Thermo, LC0725). Electrophoresis (4°C, 100 V then 15 mA) used cathode buffer B (50 mM tricine, 7.5 mM imidazole, 0.02% G-250), switched to buffer B/10 (0.002% G-250) at one-third migration and run to the bottom (∼3–4 h). For complexome profiling, each lane was sliced into 60 equal fractions. (Blue native PAGE was also used for the ATP5I dimer immunoblots in Figure 4J; transfer is described under western blotting.)

#### In-gel tryptic digestion

Gel slices were destained (50% ACN, 50 mM ammonium bicarbonate), reduced/alkylated (TCEP 100 mM, 2-chloroacetamide 400 mM), dehydrated with ACN, and rehydrated with trypsin (∼16 ng/µL) in 25 mM ammonium bicarbonate for overnight digestion at room temperature. Peptides were extracted (50% ACN, 5% formic acid), dried, resuspended in 0.2% TFA, and desalted on a Peptide Clean-up Plate (Thermo, A57865), then resuspended in 0.2% formic acid.

#### Liquid chromatography and mass spectrometry

Peptides were separated on a Vanquish Neo UHPLC (12.8 min gradient) on a µPAC HT Neo column (75 µm × 5.5 cm, 50°C) at 1.5 µL/min. Data were acquired on an Orbitrap Astral in DIA mode (MS1 Orbitrap 240,000, m/z 380–980, AGC 3.0 × 10⁶, 5 ms; DIA Astral, 2 Th windows, 299 events, HCD 26%, AGC 5.0 × 10⁴, 3 ms, 0.6 s cycle). Samples were run from lowest to highest MW slice (1–60).

### Affinity enrichment proteomics

#### Crosslinking

Crude mitochondria (500 µg) were crosslinked with disuccinimidyl sulfoxide (DSSO; Thermo, A33545). Pellets were resuspended in HEENK buffer (10 mM HEPES pH 7.1, 1 mM EDTA, 1 mM EGTA, 10 mM NaCl, 150 mM KCl) and crosslinked with 0.5 mM DSSO for 1 h at room temperature. The reaction was quenched with 100 mM Tris pH 8.0, and mitochondria were washed with HEENK/100 mM Tris.

#### MIC60 immunoprecipitation

Crosslinked mitochondria were solubilized in 50 µL solubilization buffer (50 mM imidazole, 500 mM 6-aminocaproic acid, 1 mM EDTA, 1× HALT) with digitonin (6 g/g) on ice for 20 min and cleared (15,000 × g, 20 min). The supernatant was adjusted to 250 µL with low-salt wash buffer (LSWB; 300 mM NaCl, 20 mM HEPES pH 7.8, 20% glycerol, 2 mM MgCl₂, 0.2 mM EDTA, 0.05% digitonin, 0.5 mM DTT) and incubated overnight at 4°C with rabbit IgG (Thermo, 02-6102) or anti-MIC60/IMMT (1.5 µg; Proteintech, 10179-1-AP). Protein A Dynabeads (Thermo, 10001D; 30 µL) were added for 2 h at 4°C, then washed twice each with LSWB, PBS + 0.05% digitonin, and PBS.

#### On-bead tryptic digestion

On-bead proteins were denatured in 2 M urea/100 mM Tris pH 8.0, reduced (5 mM DTT, 56°C), alkylated (15 mM iodoacetamide), and digested overnight at 37°C with 1 µg trypsin. Peptides were acidified with TFA, desalted (Thermo, 89851), dried, and resuspended in 0.2% formic acid.

#### Cell growth assay

Cells were seeded into 96-well imaging plates (Corning) at 10,000 per well (three wells per condition) and imaged in an Incucyte S3 at 37°C/5% CO₂ with a 10× objective every 1.5 h for 24 h; proliferation was quantified as phase-object count per mm². The specific growth-rate constant µ was the slope of ln(count) versus time over 3–15 h; doubling time was ln(2)/µ. Per-well µ were compared by one-way ANOVA with Tukey’s HSD (mean ± SEM; significant at P < 0.05).

#### Oxygen consumption rate

OCR was measured on a Seahorse XF96 (Agilent). AML12 cells (30,000/well) or HAP1 cells (40,000/well) were seeded in XFe96 microplates 24 h prior. Medium was replaced with XF DMEM (pH 7.4; Agilent, 103575-100) with 10 mM glucose or galactose and 2 mM L-glutamine, and equilibrated 1 h at 37°C without CO₂. The Mito Stress Test (Agilent, 103015-100) sequentially injected oligomycin (AML12 1.5 µM; HAP1 2.0 µM), FCCP (AML12 1 µM; HAP1 0.5 µM), and rotenone/antimycin A (0.5 µM each). OCR was normalized to DNA (Quant-iT PicoGreen; Thermo, P7589).

## QUANTIFICATION AND STATISTICAL ANALYSIS

### Proteomics data analysis

#### DDA data, Orbitrap Exploris 240

DDA data were analyzed in Proteome Discoverer (v2.5.0.400) with SequestHT against a mouse reference proteome (UniProt SwissProt canonical, taxonomy 10090, downloaded 2022-06-14) plus contaminants, full tryptic specificity, ≤2 missed cleavages, peptide length 6–30. Precursor/fragment tolerances were 10 ppm/0.02 Da. Fixed: carbamidomethyl-Cys. Variable: Met oxidation, phospho-STY, (≤6 per peptide). SILAC channels: medium Lys4/Arg6, heavy Lys8/Arg10. PSMs were validated by Percolator (1%/5% FDR). Phosphosite localization used IMP-ptmRS (≥75% probability). Features were aligned with Minora. SILAC quantification was at MS1 (top-3), with background-based t-tests. Proteins were annotated against MitoCarta3.0. Total-proteome samples omitted phospho and used ≤3 modifications.

#### DIA data, Orbitrap Astral

DIA files were processed in Proteome Discoverer (v3.2.0.450) using CHIMERYS (v4.0.20; INFERYS 4.7.0) via Ardia, against mouse and human reference proteomes where appropriate (taxonomy 10090, Release 406, 2024-10-02; taxonomy 9606, Release 2026-01-09) plus contaminants, Trypsin/P, ≤3 missed cleavages, length 7–30, charge 1–4, fragment tolerance 20 ppm. Fixed: carbamidomethyl-Cys. Variable: Met oxidation, phospho-STY (≤4 per peptide). Validation at 1%/5% FDR; quantification by MS2 apex with Quan in All Files. Consensus processing included strict-parsimony grouping and phosphosite annotation. Total-proteome samples omitted enrichment and phospho.

#### Phosphoproteomic and proteomic data processing

Quantitative data were generated for ten phosphatase-knockdown targets (HDHD5, PDP1, PGAM5, PPM1K, PPM1M, PPTC7, PTPMT1, PTPN11, PTPN21, PTPN4) versus non-targeting control, four replicates per channel. Processing used R (v4.5.2) with tidyverse, data.table, readxl, here, and ggpubr. Phosphopeptides were filtered, razor peptides collapsed, and contaminants removed. Intensities were median-normalized in log2 space; missing values were imputed (1–2/4 from the row’s mean/SD; 3–4/4 from a low-abundance distribution). Each phosphopeptide’s abundance was protein-normalized by subtracting the mean parent-protein log2 abundance. Per target, protein-normalized abundance was compared by two-sided Student’s t-test with Benjamini–Hochberg correction. A phosphopeptide was dynamic when |log2FC_normProt| ≥ 1 and adjusted p < 0.05. The dataset comprised 11,678 unique phosphopeptides (415 mitochondrial; 165 dynamic), corresponding to 350 localized mitochondrial phosphosites on 190 proteins (157 dynamic). Rescue proteomes (PGAM5, MIC19, ATP5I) were log2/median-normalized, imputed from a left-shifted Gaussian, and tested by Welch t-test with BH FDR (|log2FC| ≥ 1, adjusted p < 0.05).

#### Solvent accessibility, disorder, and secondary structure of phosphosites

Structural analyses used the confidently localized mitochondrial phosphosites as foreground (349 mapped to AlphaFold models; one excluded for a sequence mismatch) and all other S/T/Y residues in the same 190 proteins as background (10,464 residues). Predicted monomer structures were retrieved from the AlphaFold Protein Structure Database (AlphaFold2; 190/190 coverage). Per-residue SASA was computed with Shrake–Rupley (Biopython; probe 1.4 Å, 100 points/atom) and converted to relative solvent accessibility (RSA) using Tien et al. (2013) maxima; RSA ≥ 0.20 was exposed. Disorder was predicted with NetSurfP-3.0 (probability > 0.5 disordered). Distributions were compared by two-sided Mann–Whitney U; logistic regression tested whether RSA/disorder distinguished dynamic from static sites (odds ratios per 1 SD). Secondary structure was read from DSSP records in the AlphaFold mmCIF files and collapsed to helix/sheet/loop.

#### Active-site proximity and catalytic-residue analysis

Enzymes were identified by EC number per UniProt accession. Catalytic and ligand-binding residues were taken from UniProt “Active site” and “Binding site” annotations and cross-referenced with InterPro/Pfam; enzymes lacking annotation were gap-filled with AlphaFill (cofactors transplanted by homology). Distances were computed with Biopython as Cα–Cα (phosphosite to each active/binding residue, minimum retained) and, for the 53 dynamically regulated enzyme sites, as the phospho-acceptor hydroxyl oxygen to the nearest active-site/ligand atom.

#### Mitochondrial phosphosite enumeration and density

The mouse mitochondrial phosphoproteome was cataloged against MitoCarta3.0 (1,140 proteins). Known phosphosites were the union of dbPTM and EPSD (Mus musculus), matched on accession + position, yielding 981 proteins with ≥1 reported phosphoisoform and 9,168 unique phosphoisoforms; functional characterization was defined by PhosphoSitePlus Regulatory Sites. Phosphosite density (sites per residue) was computed for all identified sites (350 on 190 proteins) and the dynamic subset (157 on 106 proteins) and summarized as histograms (bins of 2.5 sites per 1,000 aa).

#### MTS-proximal phosphorylation analysis

Presequences and MPP cleavage sites were predicted with MitoFates v1.2 (metazoa). Phosphorylation was quantified as unique phosphosites per 100 S/T/Y, pooled across 563 MTS-positive proteins, in an 11-residue window from −20 to +100 relative to the cleavage site. Sites were grouped into MTS (≤0), proximal (+1 to +30), and distal (>30); enrichment of dynamic phosphorylation in proximal versus distal was tested by two-sided Fisher’s exact test.

#### Branched-chain amino acid quantification

Compounds were quantified in TraceFinder 5.1 (5 ppm) with manual peak review. Arginine (outside BCAA catabolism, low CV, detected in all samples) was the internal standard. Normalized value = (heavy-BCAA peak area) / (total protein) / (arginine peak area) × 10¹² (scaling constant). The labeled-pool change over an interval was Δ = mean_t2_ − mean_t1_.

#### Phosphosite-weighted MitoCarta pathway enrichment

The background was the detected mitochondrial proteome (828 MitoCarta3.0 proteins quantified across the ten per-line runs). Proteins were mapped to MitoPathways (≥3 background proteins tested). Each protein was weighted by its number of dynamic phosphosites, and each pathway’s member proteins were compared to the remainder by one-sided Mann–Whitney U with BH FDR.

#### Analysis of complexome profiling data

Complexome data from two replicates each of AML12 control and PGAM5-knockdown mitochondria were extracted from Proteome Discoverer, gene symbols de-duplicated (highest unique peptides, then PSMs), and missing values zero-filled. Abundances were normalized per fraction (to fraction total) then per protein (to protein maximum, 0–1). Fraction molecular weight was estimated from NativeMark markers by an exponential-decay model (MW = A × exp(B × slice)). Only proteins in all replicates were retained; migration was visualized as heatmaps and line-density plots.

## ACKNOWLEDGEMENTS

We would like to thank the Pagliarini Lab for their helpful feedback and discussion throughout the duration of this study, Chelsea Hackbart for technical assistance, Juan Liu for assistance with mass spectrometry method development, Natalie Niemi and Paul Grimsrud for feedback and helpful insights, the Washington University Diabetes Research Center (NIH 5P30 DK020579) for use of instrumentation, and the Washington University Center for Cellular Imaging (WUCCI; supported by Washington University School of Medicine, The Children’s Discovery Institute of University and St. Louis Children’s Hospital (CDI-CORE-2015-505 and CDI-CORE-2019-813) and the Foundation for Barnes-Jewish Hospital (3770 and 4642)). This work was supported by NIH awards R01 DK098672 and R35 GM131795 (D.J.P.), as well as funds from the BJC Investigator Program (to D.J.P.). This work was supported by the European Molecular Biology Organization (ALTF 263-2022) and the Swiss National Science Foundation (P500PB_211038) (both to P.F.). D.J.P. is an investigator of the Howard Hughes Medical Institute. This article is subject to HHMI’s Open Access to Publications policy. HHMI lab heads have previously granted a non-exclusive CC BY 4.0 license to the public and a sublicensable license to HHMI in their research articles. Pursuant to those licenses, the author-accepted manuscript of this article can be made freely available under a CC BY 4.0 license immediately upon publication.

## AUTHOR CONTRIBUTIONS

Conceptualization: A.J.S., D.J.P. Data curation: A.J.S. Formal analysis: A.J.S., P.F. Funding acquisition: D.J.P. Investigation: A.J.S., P.F., S.R., M.F. Methodology: A.J.S. Project administration: A.J.S. Resources: D.J.P. Supervision: A.J.S., D.J.P. Validation: A.J.S. Visualization: A.J.S., P.F. Writing – original draft: A.J.S., D.J.P. Writing – review & editing: A.J.S., D.J.P. All authors critically reviewed and approved the final version of the manuscript.

## DATA AND CODE AVAILABILITY

All data produced in the present study are available upon reasonable request to the authors. All raw mass spectra files supporting the findings of this study are available on the PRIDE database^57^ under the accession numbers: PXD084235, PXD084307, PXD084349, PXD084375.

## DECLARATION OF AI-ASSISTED TECHNOLOGIES IN THE WRITING PROCESS

During the preparation of this work the authors used the Claude platform to assist in proof reading and light editing. After using this service, the authors reviewed and edited the content as needed and take full responsibility for the content of the published article.

## SUPPLEMENTAL INFORMATION

All data produced in the present study are available upon reasonable request to the authors.

- Table S1. Phosphoproteomic and proteomic datasets of phosphatase CRISPRi experiment, related to Figs. 1, 2, 3, and 4.
- Table S2. Phosphoproteomic and proteomic datasets of PGAM5 rescue experiment, related to Fig. 4.
- Table S3. Proteomic datasets of MIC19 and ATP5I rescue cell lines, related to Fig S4.
- Table S4. Complexome profiling dataset of control and PGAM5-KD mitochondria, related to Fig. 5.
- Table S5. Proteomic dataset of MIC60 affinity enrichment experiment, related to Fig. 5.

---

**Figure S1.**
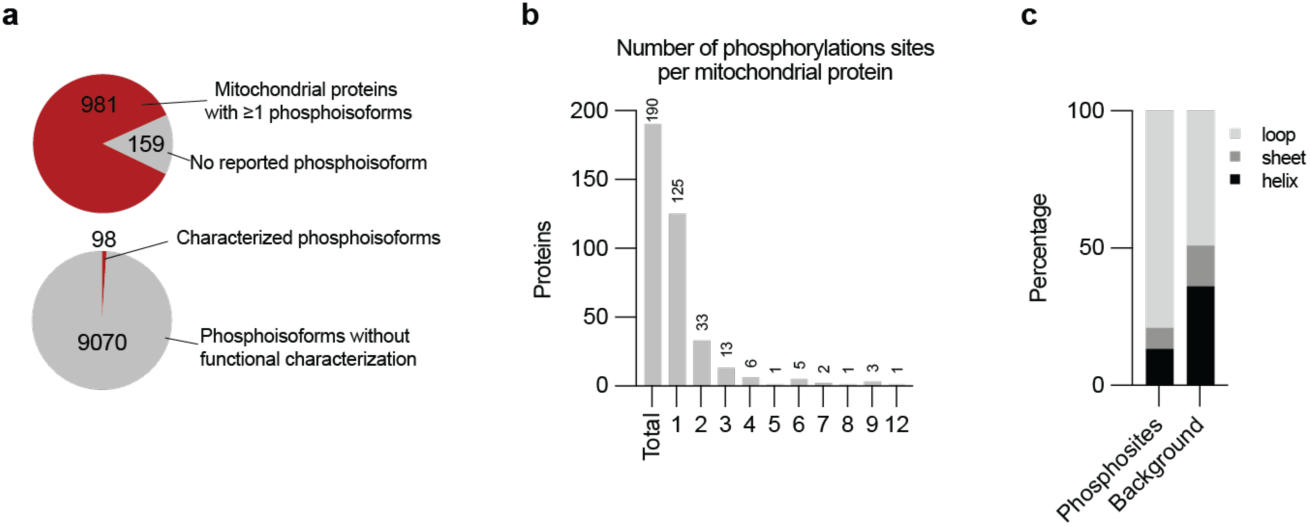
(A) Fraction of MitoCarta3.0 proteins with at least one detected phosphoisoform (981) versus none (159) (top), and fraction of detected phosphoisoforms that are functionally characterized (98) versus uncharacterized (9,070) (bottom). Characterized sites curated from PhosphoSitePlus, mapped onto the MitoCarta3.0 mouse mitochondrial proteome. Known phosphosites from dbPTM (experimental catalog filtered to MitoCarta accessions) and EPSD (Mus musculus). (B) Number of phosphorylation sites per mitochondrial protein (n = 190 phosphorylated proteins; median = 1). (C) Secondary-structure composition (loop, sheet, helix) of phosphosites versus all other S/T/Y residues in the same proteins as background.

**Figure S2.**
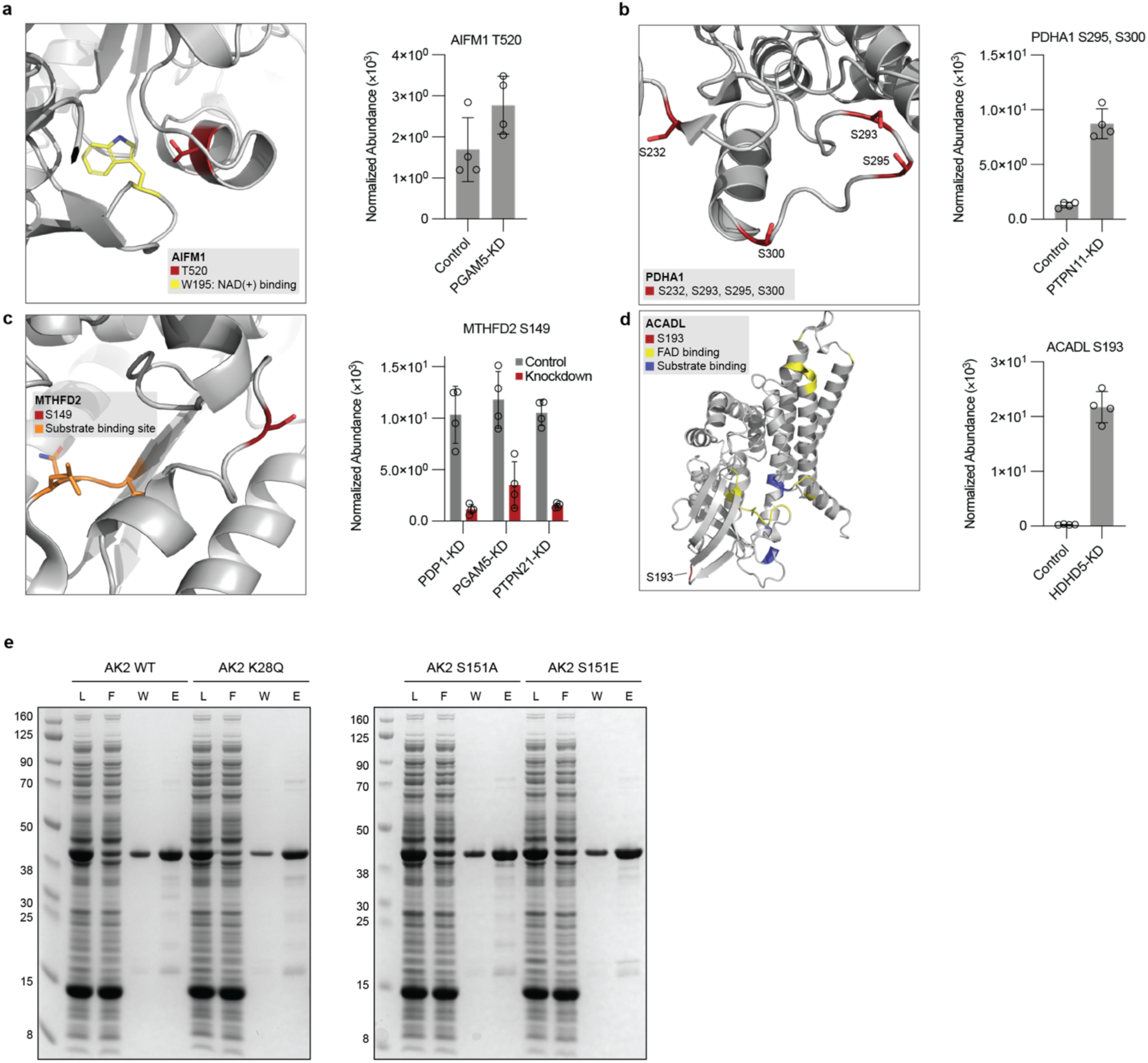
(A) T520 in proximity to W195 in AIFM1, with protein normalized phosphopeptides abundance in PGAM5-KD. (B) PDHA1 S232, S293, S295, and S300 mapped onto protein structure. PDHA1 S295, S300 protein-normalized phosphopeptide abundance in PTPN11-KD. (C) S149 of MTHFD2 mapped in proximity to substrate binding site residues. MTHFD2 S149 protein-normalized phosphopeptide abundance in control and knockdown lines. (D) ACADL S193 in proximity to FAD and substrate binding sites. ACADL S193 protein-normalized phosphopeptide abundance in HDHD5-KD cells. (E) Coomassie stained SDS-PAGE gel of purified recombinant SUMO-tagged AK2 (WT, K28Q, S151A, S151E). L = loading, F = flow-through, W = wash, E = elution.

**Fig. S3.**
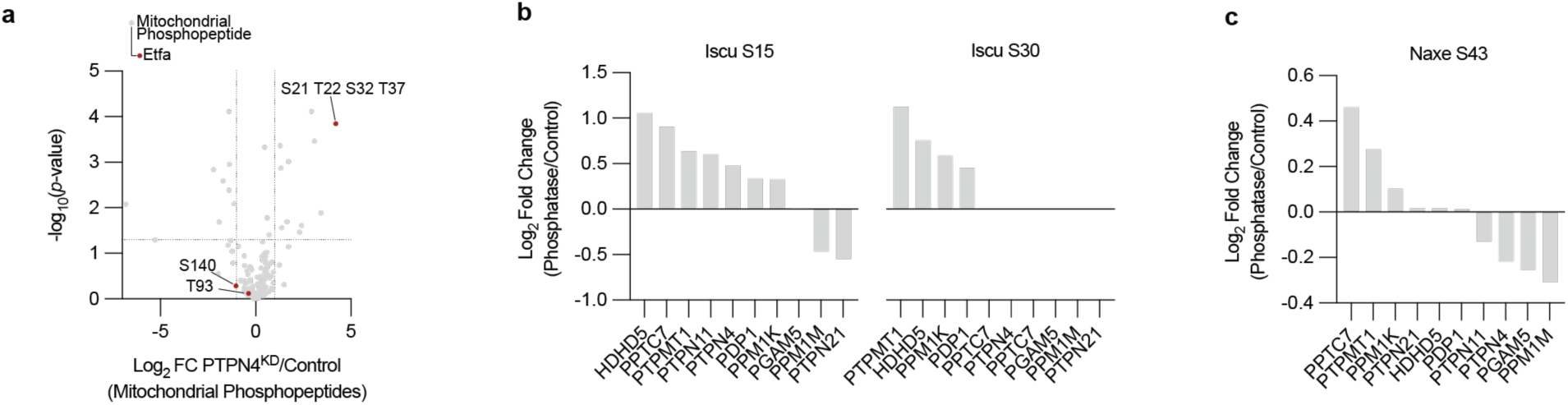
(A) Volcano plot of mitochondrial phosphopeptides in PTPN4-KD/Control cells showing significant elevation for a 4x phosphorylated Etfa peptide relative to two unchanged Etfa phosphopeptides. (B) Iscu S15, and S30 log2 fold change of phosphatase knockdown lines compared to matched controls. (C) Naxe S43 log2 fold change of phosphatase knockdown lines compared to matched controls.

**Figure S4.**
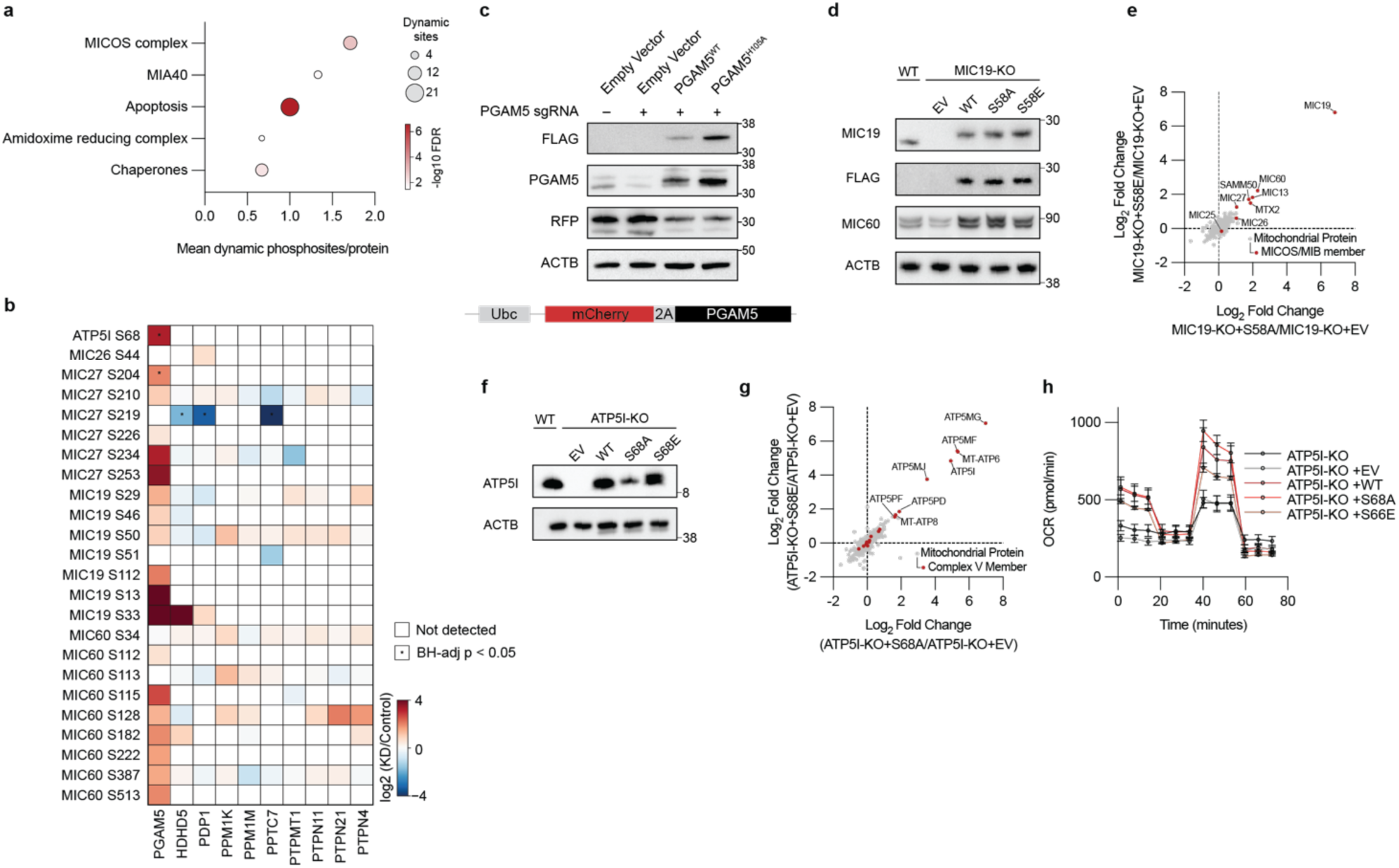
(A) Phosphosite-weighted over-representation of MitoCarta pathways: each of the 828 detected mitochondrial proteins (union across the 10 per-line proteomes) is weighted by its number of dynamically regulated phosphosites (0 if none); pathway members compared to the rest of the proteome by one-sided Mann-Whitney U test, BH FDR. (B) Log2 FC (KD/control) of all 23 quantified cristae formation phosphosites (ATP5I, MIC60, MIC19, MIC25, MIC27, MIC26) across all 10 phosphatase-knockdown lines. (Welch t-test, n = 4 SILAC replicates per line). (C) Rescue construct schematic (ubiquitin promoter–mCherry-2A-PGAM5) and immunoblots confirming PGAM5 knockdown and WT/H105A re-expression (PGAM5 and C-terminal FLAG). (D) Immunoblots of MIC19-KO cells reconstituted with WT, S58A, or S58E (MIC19, FLAG, MIC60, ACTB). (E) Proteomic scatter, MIC19-KO+S58A/ vs MIC19-KO+S58E (log2 fold change over MIC19-KO+EV). Welch t-test, BH adjusted p-value; n = 3. (F) Immunoblots of ATP5I-KO cells reconstituted with WT, S68A, or S68E (ATP5I, ACTB). (G) Proteomic scatter, ATP5I-KO+S68A/EV vs ATP5I-KO+S68E/EV; defects restored by both. Welch t-test, BH adjusted p-value; n = 3. (H) Seahorse OCR (pmol/min) of ATP5I-KO and rescues (EV, WT, S68A, S68E) in glucose.

**Figure S5.**
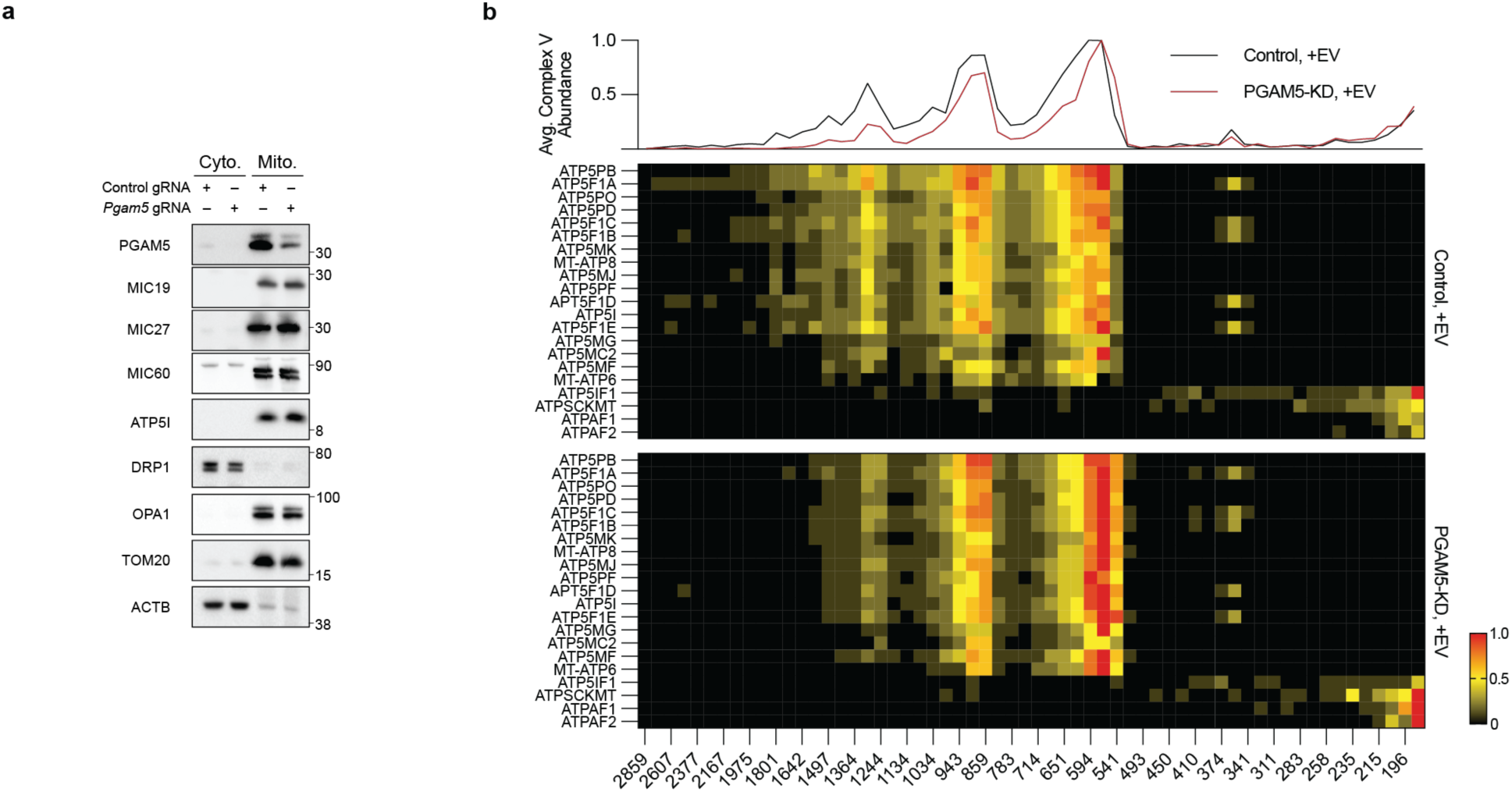
(A) Immunoblots of purified mitochondrial and cytosolic fractions (control, PGAM5-KD) for PGAM5, MIC19, MIC27, MIC60, ATP5I, DRP1, OPA1. TOM20 and ACTB were used as mitochondrial and cytosolic loading controls. (B) Complexome profiling heatmap and line profile of complex V (control vs PGAM5-KD); reduced dimeric/oligomeric-to-monomeric ratio in PGAM5-KD.

